# Environmental Contributions to Proton Sharing in Protein Low-Barrier Hydrogen Bonds

**DOI:** 10.1101/2025.11.05.686872

**Authors:** Jiusheng Lin, Oksana Gerlits, Daniel W. Kneller, Kevin L. Weiss, Leighton Coates, Mark A. Hix, Solomon Y. Effah, Andrey Kovalevsky, Alice R. Walker, Mark A. Wilson

## Abstract

Hydrogen bonds (H-bonds) are central to biomolecular structure and dynamics. Although H-bonds are typically characterized by well-defined proton positions, proton delocalization can play a key role in facilitating enzyme catalysis and allostery in some systems. Experimentally locating protons is difficult, hampering the study of the proton mobility in H-bonds. We used neutron crystallography and large quantum mechanics/molecular mechanics-Born Oppenheimer molecular dynamics (QM/MM-BOMD) simulations of human DJ-1 and its bacterial homolog YajL to validate atomic resolution X-ray crystallographic bond length analysis, directly visualizing the shared deuteron in a low-barrier hydrogen bond (LBHB) between Glu14-Asp23 in YajL that is a conventional H-bond in DJ-1. In addition, X-ray bond length analysis of protiated and perdeuterated DJ-1 and YajL shows no significant effect of deuteron substitution on these carboxylic acid-carboxylate H-bonds but does reveal an effect at the active site glutamic acid. Residues in the vicinity of Glu14-Asp23 that might favor LBHB formation in YajL were interrogated by mutagenesis of homologous residues in DJ-1. X-ray bond length analysis and QM/MM-BOMD simulations demonstrate that a distal I21T DJ-1 substitution increases proton delocalization in the Glu-Asp H-bond. In addition, simulations show that the extent of proton mobility in the H-bond influences correlated dimer-spanning motions in YajL and DJ-1. In total, we show that mutations within extended H-bonding networks can modulate proton transfer barriers in carboxylic acid-carboxylate H-bonds, allowing proton delocalization to be engineered using combined bioinformatic, structural, and computational information.

## Introduction

Low-barrier hydrogen bonds (LBHBs) are H-bonds where the barrier to proton transfer from the donor to the acceptor atom is comparable to the quantum mechanical zero-point energy of the interaction (*1, 2*) (Figure 1A,B). Facile proton transfer makes LBHBs effective contributors to acid-base catalysis in certain enzymes (*2-7*), creating favored pathways of allosteric communication between distant sites during catalysis (*8*), and imparting binding selectivity and specific elicited responses with certain substrates (*9-12*). Because of their utility in enhancing various aspects of protein and enzyme function, understanding which features of an H-bond are important for lowering the proton transfer barrier and creating an LBHB would be useful for the prediction and engineering of dynamic hydrogen bonded networks.

**Figure 1:**
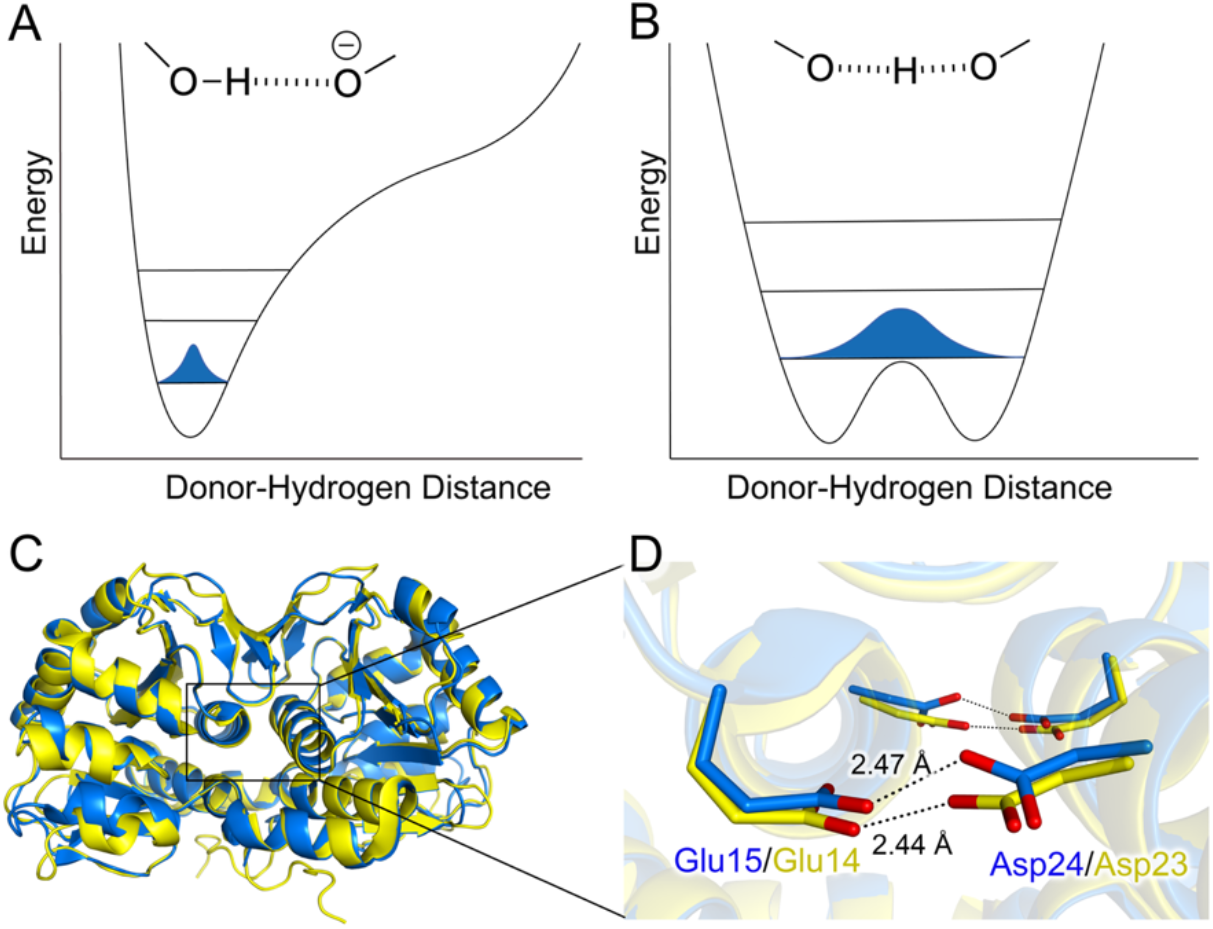
Conventional H-bonds and LBHBs in DJ-1 and YajL. (A) An energy diagram of a conventional OH-^-^O H-bond is shown, with the vibrational levels shown as horizontal lines and the probability of proton location (corresponding to the quantum probability density function Ψ^2^) shown in blue curve. (B) shows an LBHB where the proton transfer barrier is slightly lower than the zero-point energy of the ground vibrational level. The highest probability for proton location (maximum of the blue curve) is in the middle of the oxygen atoms. (C) Dimeric human DJ-1 (blue) and YajL (yellow) are superimposed, revealing their highly similar backbone structures. (D) is an expanded region of the DJ-1 and YajL dimer, with residues Glu15 (blue, DJ-1)/Glu14 (yellow, YajL) and Asp24 (blue, DJ-1)/Asp23 (yellow, YajL) shown. The short H-bonds are indicated with dotted lines.

Despite their importance in biochemistry, reliably identifying LBHBs presents challenges. Experimental methods for determining LBHBs include neutron crystallography (*13-16*), atomic resolution (d_min_<1.2 Å) X-ray crystallography (*17-19*), NMR spectroscopy (*20, 21*), IR spectroscopy (*22, 23*), and H/D exchange kinetics (*24*), each of which has limitations. Of these methods, only crystallographic approaches can determine the H atom position in the bond. Neutron crystallography permits unambiguous determination of hydrogen position but requires large crystals that can be challenging to grow, as well as access to limited neutron beamtime. Atomic resolution X-ray crystallography can sometimes locate hydrogen; however, the small X-ray form factor of hydrogen limits the signal generated from hydrogen atoms, allowing only the location of electrons to be reported, not protons. These effects combine to diminish both the precision and accuracy of hydrogen atom location in atomic resolution electron density maps produced by X-ray crystallography (*25*).

Bond length analysis using atomic resolution X-ray crystallography can also be used to infer the presence of H atoms on certain residues such as carboxylic acids owing to protonation effects on bond lengths between the more electron-rich carbon and oxygen atoms, even when electron density for the hydrogen atom cannot be observed. Determining protonation states using bond length analysis relies on the precise measurement of differences in C-O bond length in protonated vs. deprotonated carboxylic acids (*17, 26-29*). In a fully protonated carboxylic acid, the C=O bond is ∼1.21 Å and the C-OH bond is ∼1.31 Å. In a deprotonated carboxylic acid, both C-O bonds have similar lengths (1.26 Å) due to resonance. Prior studies applied bond length analysis to human DJ-1 and *E. coli* YajL, two homodimeric proteins of the DJ-1 superfamily that share 41% sequence identity and have highly similar structures (1.1 Å C*α* RMSD) (*18*). A conserved 2.4-2.5 Å hydrogen bond between Glu15 and Asp24 (DJ-1 numbering) spans the dimer and increases protein stability (*18*)(Figure 1C,D). This interaction was a candidate for a low-barrier hydrogen bond owing to its short length, and prior analysis of the Asp/Glu C-O bond lengths in atomic resolution X-ray crystal structures of these two proteins indicated there were different apparent degrees of proton delocalization in the Glu-Asp H-bond, with YajL potentially harboring a LBHB (*18*). This result has also been very recently supported by new quantum mechanics-based X-ray refinement methods (*30*). Despite its potential utility, X-ray bond length analysis for LBHB identification has not been validated by comparison to neutron diffraction data, which can directly visualize protons and deuterons.

In part because of the difficulty in reliably identifying LBHBs, the structural determinants of LBHBs are not well understood. The free energies of the protonated donor and acceptor atoms in an LBHB must be comparable for the proton transfer barrier to be low enough to permit transfer (Figure 1B), which is equivalent to requiring that pK_a_ values of donor and acceptor in LBHBs be similar (*3*). Therefore, protein microenvironmental features that alter pK_a_ values should be important for creating permissive environments for LBHBs. Though several different types of LBHB have been identified in proteins (*31*), hydrogen bonds between carboxylic acid-containing amino acids (Asp and Glu) have attracted special interest, as they may frequently meet the matching pK_a_ requirement (*18, 32-34*). In addition, the short donor-acceptor oxygen distance of ∼2.5 Å in Asp/Glu hydrogen bonds makes them frequent suspects for LBHBs, although prior work has shown that some of these hydrogen bonds have largely localized protons participating in standard hydrogen bonds (*18, 34, 35*).

Because the zero-point energy of the LBHB must be close to the proton transfer barrier height, quantum mechanical treatments are essential for their computational characterization (*1, 36-38*). Advances in QM/MM simulations have increased the size of the region that can be treated quantum mechanically including dynamics, moving from a few atoms to 100s, encompassing several amino acids. These improvements can more fully incorporate quantum mechanical environmental influences on the proton transfer barrier, permitting the direct simulation of proton transfer without the use of biasing potentials or other non-physical computational interventions. Moreover, proteins are dynamic molecules that sample an ensemble of nearly isoenergetic conformations under physiological conditions (*39*). As the hydrogen-bonding microenvironment fluctuates in the ensemble, the proton transfer barrier becomes multidimensional and time-dependent (*40*). Motional modes that couple strongly to the proton transfer reaction coordinate will influence the degree of proton sharing and may lead to switching between conventional and LBHB character (*41*). Large QM-region, dynamic QM/MM-MD simulations offer the promise of providing an integrated electronic and dynamic picture of LBHBs that can more fully explain their essential structural determinants. In addition, new quantum mechanics-based X-ray refinement methods can identify LBHBs from X-ray diffraction data at moderate resolutions (*30*), illustrating the power of combining more accurate QM physical models and X-ray experimental data to produce models with better representations of uncommon types of interaction.

In this work, we use a combination of neutron crystallography, atomic resolution X-ray crystallography, large quantum mechanics/molecular mechanics (QM/MM) simulations, and mutagenesis to investigate environmental effects in these homologous proteins that influence proton delocalization in a conserved H-bond. Our results indicate that X-ray bond length analysis is a reliable tool for identifying carboxylic acid-carboxylate LBHBs and that new QM/MM methods combined with homology-informed mutations permit engineering of H-bonds networks that enhance proton delocalization.

## Results

### Neutron crystal structures and QM/MM simulation validate X-ray bond length-based identification of a low-barrier hydrogen bond in YajL

Prior atomic resolution X-ray bond length analysis unexpectedly indicated that DJ-1 has a conventional short H-bond between Glu15 and Asp24 while YajL may have a LBHB at the same location (Glu14/Asp23), although the LBHB assignment in YajL was tentative (*18*). To establish the status of these two structurally similar H-bonds, we used neutron crystallography to directly visualize the location of the hydrogen (deuterium) atom in this interaction. Perdeuterated DJ-1 and YajL proteins were produced at Oak Ridge National Laboratory (ORNL) (see Methods) and used to grow large (∼2×1×0.5 mm^3^) crystals (Figure S1) for neutron diffraction, which was performed at the MaNDI instrument at the Spallation Neutron Source, ORNL. In both DJ-1 and YajL nuclear density maps, 3σ positive difference peaks were observed between the oxygen atoms of the H-bonded carboxylic acid residues, consistent with the presence of a deuteron (Figure 2). Because deuterium atoms scatter neutrons strongly (coherent cross section of ^2^H for thermal neutrons is 5.6 barn, the same as for ^12^C) (*42*), the locations of these deuterons can be determined from moderate resolution neutron nuclear density maps. In the 2.15 Å resolution nuclear density map for human DJ-1, the deuterium peak is within ∼0.8 Å of Asp24 (Figure 2). This O-D distance is ∼0.2 Å shorter than expected but indicates that DJ-1 has a predominantly localized proton on Asp24 in a conventional Asp24-Glu15 H-bond. The quantitative disagreement between the nuclear density centroid and ideal O-D distance may be owing to the unrestrained refinement of the D atom position to avoid restraint bias and to the 2.15 Å resolution of the data. By contrast, the 1.79 Å resolution nuclear density map for YajL shows this deuteron density peak positioned nearly equidistant between Glu14 Oε2 and Asp23 Oδ2 in both protomers (Figure 2). These neutron results are consistent with the prior X-ray bond length analysis of protiated DJ-1 and YajL (*18*) and quantitatively similar to recent quantum mechanical refinement results (*30*), confirming that DJ-1 has a conventional COOH-^-^ OOC H-bond while YajL has a low-barrier hydrogen bond with the deuteron shared between Asp23 and Glu14.

**Figure 2:**
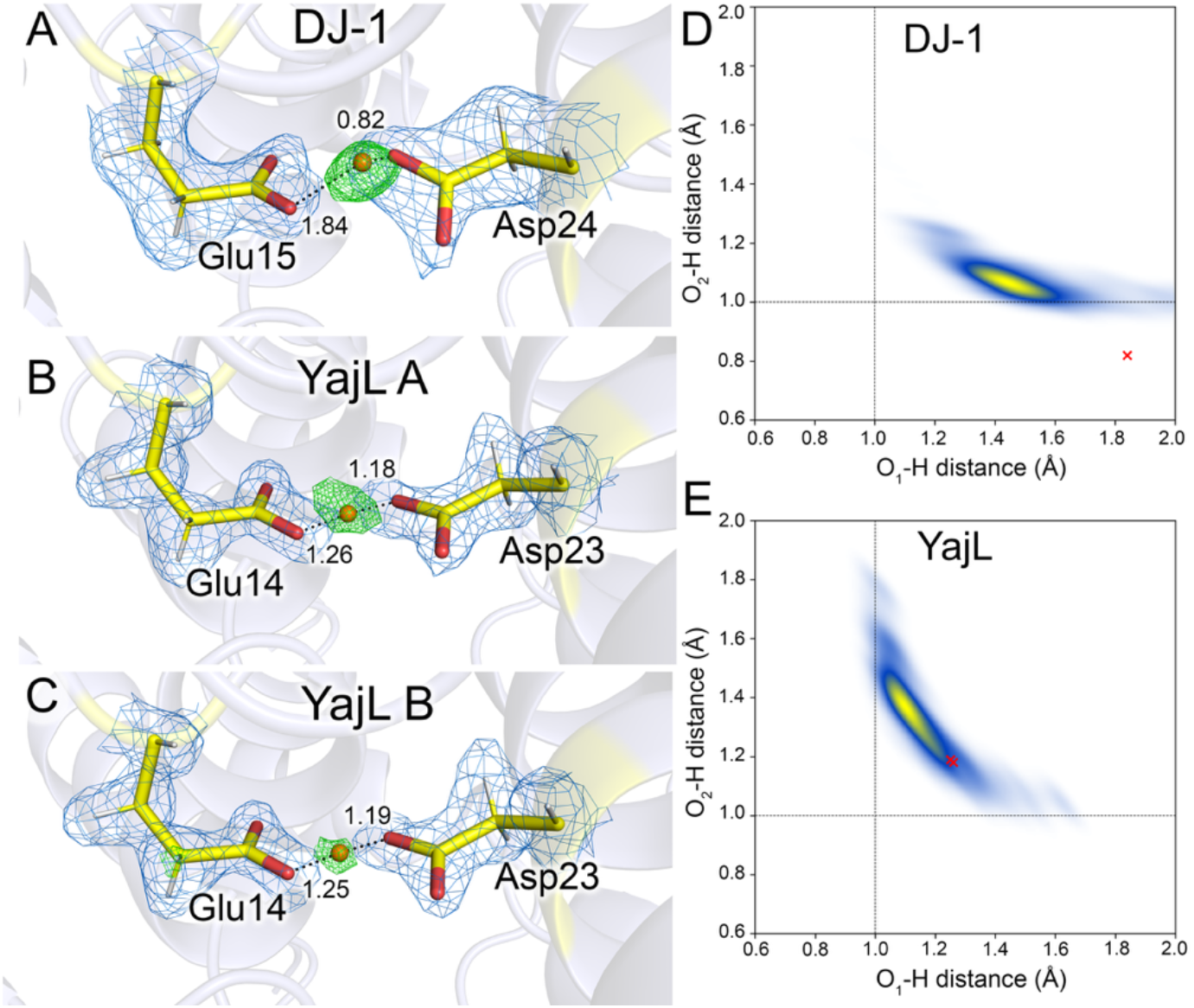
Degree of proton/deuteron sharing in the short H-bond in DJ-1 and YajL. Panels A-C show nuclear density maps calculated from neutron diffraction data, with 2mF_o_ – DF_c_ nuclear density (blue) contoured at 1.2*σ* and the mF_o_ − DF_c_ nuclear density contoured at 2.9-3.3*σ* (positive, green; negative, red). In the 2.15 Å resolution map for DJ-1 (A), the difference peak for the deuteron in the Glu15−Asp24 H-bond indicates that the hydrogen is associated with Asp24. Panels B and C show the 1.79 Å resolution maps of YajL for the two independent Glu14−Asp23 H-bonds in the asymmetric unit. (YajL A/B). The deuteron in the mF_o_ − DF_c_ difference nuclear density peak is more symmetrically positioned in YajL and indicates an LBHB. Panels D and E show the QM/MM-BOMD simulation-derived density of states for the position of the proton in the Glu-Asp H-bond in DJ-1 (D) and YajL (E), with the red crosses indicating the experimentally determined unrestrained refined location of the deuterons from neutron diffraction. In DJ-1 (panel D), simulation shows a hydrogen atom localized predominantly on Asp24 (corresponding to the O_2_H distance on the Y-axis). By contrast, in YajL (panel E) the hydrogen is localized more symmetrically between the oxygen atoms of Glu14 (O_1_) and Asp23 (O_2_), consistent with an LBHB.

Quantum mechanics/molecular mechanics with Born-Oppenheimer molecular dynamics (QM/MM-BOMD) simulations provide detailed and easily interpretable information about proton behavior in a candidate hydrogen bond. Sampling the proton’s vibrations over time allows the degree of localization of the proton within the hydrogen bond to be quantified. A typical oxygen-containing H-bond will have a donor O-H distance of ∼1 Å and an acceptor -H^…^O distance of ∼1.7 Å. In conventional short, charge-assisted COOH-^-^OOC H-bonds, the total donor-acceptor O-O distance is 2.4-2.5 Å, but the proton will remain localized to the donor atom. This is the scenario we observe in QM/MM-BOMD simulation of wild-type DJ-1 (Figure 2D), where the proton remains localized to the Asp24 Oδ2 H-donor with a distance of 1.0-1.1 Å and maintains a distance to the Glu15 Oε 2 acceptor of 1.4-1.8 Å over the simulation time (Figure 2D), in agreement with experiment. The simulations of YajL, by contrast, show an immediate shift in proton location to 1.2-1.3 Å from both the Asp23 Oδ2 donor and Glu14 Oε2 acceptor atoms, with the proton sampling the full span of possible donor/acceptor distances from 1.0-1.7 Å over time. This is indicative of an almost barrierless proton transition and provides clear support for an LBHB, since the proton is shared between the donor and acceptor (Figure 2E). The QM/MM-BOMD values agree well with the experimental results for the acceptor distances, though the centroid of the donor values is shifted by about −0.2 Å compared to the location of the deuteron in the neutron crystal structures. O-D bond lengths are approximately 0.03 Å shorter than O-H distances (*43*), although the 0.2 Å discrepancy is in the opposite direction of this effect. Relaxation of the structure in the simulation to a conformation that is subtly different than that observed in the crystal structure or errors associated with our use of the centroid of the nuclear density mF_o_-DFc peak without restraints as the deuteron location may also contribute.

LBHBs have been postulated to be stronger than conventional H-bonds (*1*), although this has been contested (*44, 45*). Having confirmed an LBHB between Glu14 and Asp23 in YajL, we tested its importance for YajL stability. The D23N YajL mutant was made and its stability assessed using differential scanning fluorimetry. This Asp->Asn mutant preserves the H-bond but eliminates its potential for proton sharing. D23N YajL is less thermally stable (T_m_=56.5°C) than wild-type (T_m_=60.5°C) (Figure 3A). This 4.0°C T_m_ decrease upon Asp->Asn mutation in YajL is larger than the 2.4°C decrease observed for the homologous D24N mutant in the conventional H-bond of DJ-1 (*18*), suggesting that the YajL LBHB is somewhat more stabilizing. Comparison of the X-ray crystal structures of wild-type (0.94 Å resolution) and D23N YajL (0.98 Å resolution) shows the expected lengthening of the H-bond from 2.44/2.46 Å (protomers A/B) in wild-type YajL to 2.79/2.80 Å (protomers A/B) in D23N YajL (Figure 3B), nearly identical to the lengthening observed in D24N DJ-1 (*18*). However, in contrast to DJ-1, the D23N YajL mutation results in disorder at the E14-N23 H-bond that was modelled with alternate confirmations in the crystallographic model (Figure 3C).

**Figure 3:**
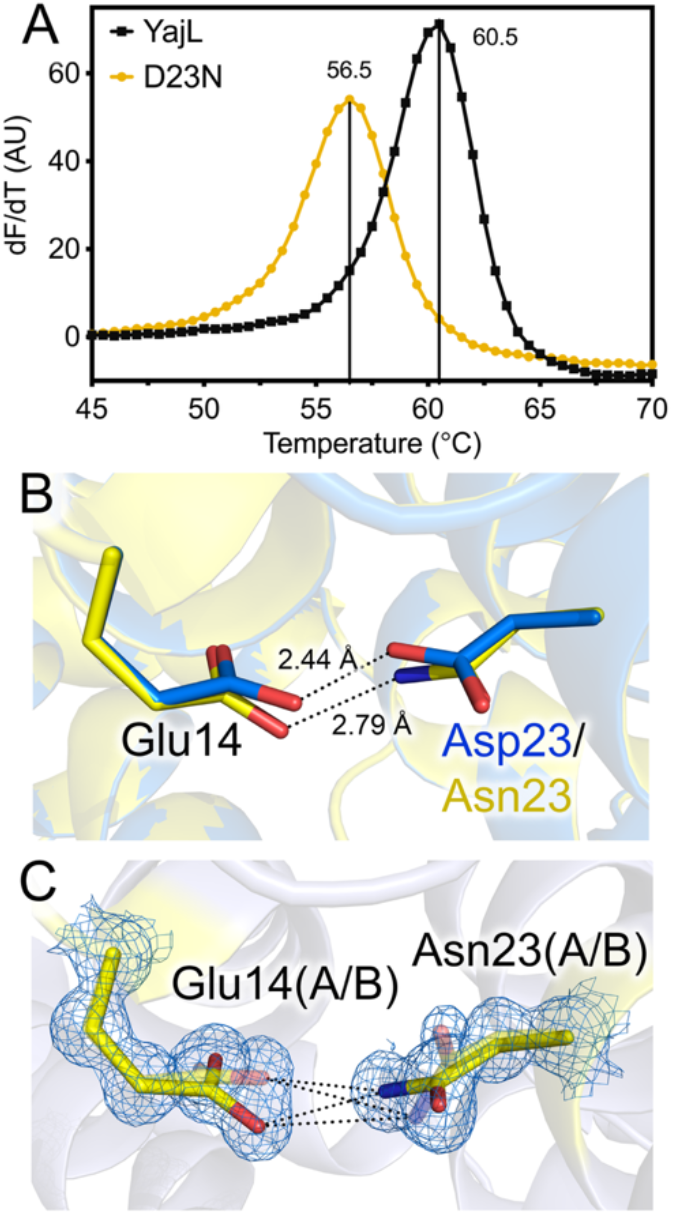
Loss of an LBHB destabilizes YajL. Panel A shows differential scanning fluorimetry (first derivative of the fluorescence signal with temperature on the Y-axis) of wild-type YajL and the D23N mutant that eliminates possibility of proton sharing in an LBHB. The conversion of the COO-H-OOC LBHB to a conventional NH-^-^OOC H-bond reduces T_m_ by 4.0 °C. Panel B show a superposition of crystal structures of wild-type (blue) and D23N YajL (yellow), confirming that the D23N mutation converts the short LBHB O-O distance to a conventional H-bond with the expected ∼2.8 Å N-O distance. Panel C shows 2mF_o_ – DF_c_ electron density for the 0.98 Å resolution D23N YajL crystal structure contoured at 1.0*σ* (blue). The conversion of the Glu-Asp LBHB to a Glu-Asn conventional H-bond results in a disordered interaction, where the minor conformations are shown with partially transparent sticks.

### Deuteration has no effect on Glu-Asp H-bond character in DJ-1 or YajL but does influence active site Glu protonation

The agreement between neutron diffraction and prior atomic resolution X-ray diffraction validates X-ray bond length analysis as a useful tool for detecting potential LBHBs between carboxylic acids. As with all macromolecular crystals used for neutron diffraction, our crystals were exchanged into D_2_O prior to data collection to avoid the large incoherent neutron scattering by hydrogen atoms. In addition, we used perdeuterated proteins to eliminate incoherent scattering from the non-exchangeable hydrogen atoms. Previous studies have found that the effect of deuterium substitution in protein crystal structures is generally minor (*46, 47*). However, it is possible that isotopic substitution effects could be larger in H-bonds. The barrier height in LBHBs is defined relative to the zero-point energy of the bond, which changes upon isotopic substitution. For this reason, LBHBs have anomalous fractionation behavior in H/D exchange (*48*) and H/D substitution alters H-bond geometries in supermolecular assemblies (*49*).

Because of the importance of deuteration for neutron crystallography and its potential influence on LBHBs, we determined the influence of D_2_O and protein perdeuteration on these Glu-Asp H-bonds by collecting atomic resolution X-ray diffraction data of protiated proteins in H_2_O and perdeuterated proteins in D_2_O. Both electron density and bond length analyses were used to interrogate the degree of protein/deuteron delocalization in the Glu-Asp H-bond. Positive mF_o_-DF_c_ difference electron density peaks at ∼3σ are observed between Glu15 and Asp24 in X-ray crystal structures of both perdeuterated and protiated DJ-1 determined at 0.92 Å and 1.05 Å resolution, respectively (Figure 4). We assign this peak to the hydrogen in this H-bond. In protiated DJ-1, the mF_o_-DF_c_ peak is located 1.76 Å from Glu15 Oε2 and 0.84 Å from Asp24 Oδ2, while it is 1.59 Å from Glu15 Oε2 and 0.95 Å from Asp24 Oδ2 in deuterated DJ-1 (Figure 4). Because hydrogen scatters X-rays weakly and the difference between electron and true nuclear position for hydrogen atoms makes accurately locating hydrogen atoms in electron density maps challenging, we also used atomic resolution X-ray bond length analysis. Estimated standard uncertainties (ESUs) for C-O bond lengths for all Asp/Glu residues were obtained using inversion of the full least squares matrix after unrestrained coordinate refinement in SHELXL (see Methods). Both protiated and deuterated DJ-1 show the same differences between C-O bond lengths in Glu15 and Asp24 with similar levels of statistical significance, indicating that the hydrogen is localized predominantly on Asp24 (Table 1). Corresponding C-O bonds in the protiated and deuterated DJ-1 Glu-Asp pairs differ by less than 2σ (Table 1).

**Table 1.** Atomic resolution X-ray bond length analysis of Asp/Glu residues.

| Protein<sup>a</sup> | Residue | C-O<sub>1</sub> (Å) | C-O<sub>2</sub> (Å) | $\Delta(\text{C-O})/(\sigma_1^2+\sigma_2^2)^{1/2}$ <sup>c</sup> | COO(H) status |
| --- | --- | --- | --- | --- | --- |
| H-DJ-1 | Glu15 | 1.235(10)<sup>b</sup> | 1.272(10) | 2.62 | deprotonated |
| H-DJ-1 | Asp24 | 1.216(9) | 1.290(9) | 5.81 | protonated |
| H-DJ-1 | Glu18 | 1.217(11) | 1.301(11) | 5.40 | protonated |
| D-DJ-1 | Glu15 | 1.255(6) | 1.274(6) | 2.24 | deprotonated |
| D-DJ-1 | Asp24 | 1.220(6) | 1.298(6) | 9.19 | protonated |
| D-DJ-1 | Glu18 | 1.239(8) | 1.277(8) | 3.36 | protonated, less than H-DJ-1 |
| H-YajL | Glu14<br>(molecule A,<br>molecule B) | 1.236(11),<br>1.220(10) | 1.305(10),<br>1.317(10) | 4.64, 6.86 | protonated,<br>protonated |
| H-YajL | Asp23<br>(molecule A,<br>molecule B) | 1.216(10),<br>1.223(11) | 1.305(10),<br>1.292(10) | 6.29, 4.64 | protonated,<br>protonated |
| H-YajL | Glu17<br>(molecule A,<br>molecule B) | 1.213(11),<br>1.211(10) | 1.305 (12),<br>1.308(11) | 5.65, 6.52 | protonated,<br>protonated |
| D-YajL | Glu14<br>(molecule A,<br>molecule B) | 1.226(8),<br>1.203(8) | 1.313(8),<br>1.302(8) | 7.69, 8.75 | protonated,<br>protonated |
| D-YajL | Asp23<br>(molecule A,<br>molecule B) | 1.225(8),<br>1.228(8) | 1.293(8),<br>1.291(8) | 6.01, 5.57 | protonated,<br>protonated |
| D-YajL | Glu17<br>(molecule A,<br>molecule B) | 1.247(8),<br>1.226(8) | 1.263(9),<br>1.305(9) | 1.33, 6.56 | deprotonated,<br>protonated |
| DJ-1 M17T | Glu15 | 1.254(8) | 1.265(8) | 0.97 | deprotonated |
| DJ-1 M17T | Asp24 | 1.196(8) | 1.285(8) | 7.87 | protonated |
| DJ-1 I21T | Glu15 | 1.244(7) | 1.295(8) | 4.80 | protonated |
| DJ-1 I21T | Asp24 | 1.213(7) | 1.298(7) | 8.59 | protonated |

<sup>a</sup>"H-" is protiated protein in H<sub>2</sub>O, "D-" is perdeuterated protein in D<sub>2</sub>O.
<sup>b</sup>parenthetical error notation is used to show estimated standard uncertainties (ESUs) from SHELXL.
<sup>c</sup> $\Delta(\text{C-O})=\text{distance}(\text{C-O}_2)-\text{distance}(\text{C-O}_1)$ ; $(\sigma_1^2+\sigma_2^2)^{1/2}$ is propagation of errors for ESUs on bond lengths.

**Figure 4:**
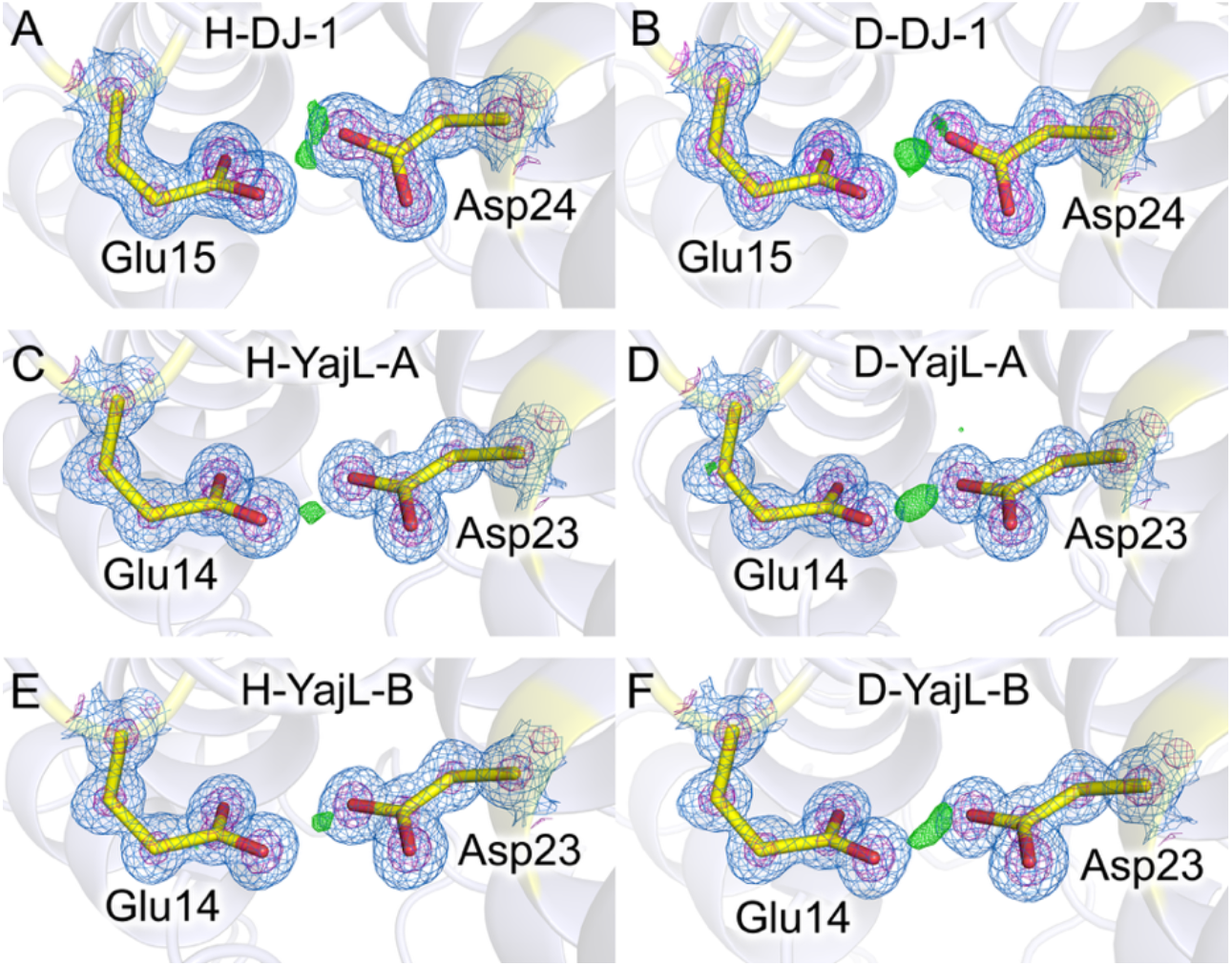
Locating the Glu-Asp H-bond hydrogen atom in atomic resolution electron density maps of protiated and deuterated DJ-1 and YajL. In all panels, 2mF_o_ – DF_c_ electron density contoured at 1.0*σ* (marine) and 5.0*σ* (purple) and mF_o_ − DF_c_ electron density contoured from 2.8-3.1*σ* (positive, green; negative, red). Protiated structures are shown in the left-hand portion (panels A,C,E) and corresponding deuterated structures are on the right (panels B,D,F). The mF_o_ − DF_c_ difference electron density peak is located between the two carboxylic acids in all panels, with a bias towards being more centrally located in YajL, consistent with an LBHB.

Like DJ-1, perdeuterated and protiated YajL (0.88 Å and 0.94 Å resolution respectively) show ∼3-4 σ peaks in mF_o_-DF_c_ difference electron density maps between Glu14 and Asp23 in both chains in the asymmetric unit (ASU) (Figure 4C-F). Unlike DJ-1, the hydrogen difference electron density peak in YajL is situated more equally between the two oxygen atoms, with the center of the mF_o_-DF_c_ peak located 1.21/1.43 Å (in the two protomers in the ASU) from Oε2 of Glu14 in protiated YajL and 1.20/1.28 Å from Oδ2 of Glu14 in the deuterated protein. The distances between the mF_o_-DF_c_ peaks and Asp24 Oδ2 atoms are also comparable between protiated and deuterated YajL (1.33/1.05 Å to Asp24 Oδ2 in protiated YajL and 1.25/1.18 Å to Asp24 Oδ2 in deuterated YajL) (Figure 4C-F). Atomic bond length analysis of protiated YajL and deuterated YajL (see Methods) confirms that both Glu14 and Asp23 are protonated at their interacting oxygen atoms (Table 1), again supporting the assignment of the Glu14-Asp23 H-bond as an LBHB (*18*). The consistent presence of an LBHB between Glu14-Asp23 demonstrates that proton delocalization in this YajL Glu-Asp H-bond is insensitive to protein or solvent deuteration. This was a point of considerable interest, as deuteration will generally reduce the zero-point energy of an H-bond and therefore might have a significant effect on LBHB character. As with DJ-1, the differences in lengths between the same C-O bond in deuterated and protiated and YajL are less than 2σ (Table 1), indicating that deuteration does not significantly alter these bond lengths or the degree of proton/deuteron localization.

To further explore the potential role of deuteration in H-bonds involving protonated carboxylic acids, we turned to the Glu18-Cys106 (DJ-1 numbering) interaction in the active sites of DJ-1 and YajL. Cys106 is the active site nucleophile and, in human DJ-1, has a pK_a_ value of 5.4 (*50*). Prior atomic resolution bond length analysis shows that the Glu is protonated, which is supported by the current analysis of both DJ-1 and YajL (*50*)(Table 1). However, bond length analysis shows that Glu18 in deuterated DJ-1 has decreased protonation (as evidenced by reduced differences between C-O bond lengths) and Glu17 in perdeuterated YajL is completely deprotonated (ionized) in one of the two protomers of YajL, while Glu17 in the other protomer appears similar to protiated YajL (Table 1). This is consistent with the absence of a strong peak for a deuteron at this location in the nuclear density maps for both DJ-1 and YajL. Furthermore, a broadly similar enrichment of a zwitterionic Cys-S^-^-His^+^ active site dyad in SARS CoV-2 main protease was observed in D_2_O compared to H_2_O (*51*). Therefore, H/D substitution has differing effects on protonation of ionizable groups and is sensitive to the microenvironment of the residue. This sensitivity seems to be enhanced in the vicinity of low pK_a_ cysteine residues, which merits further investigation.

### Proton delocalization in hydrogen bonds is affected by local environment

The sensitivity of proton location in LBHBs to the details of their environment has long been appreciated in small-molecule crystallography, where counterion identity or crystal lattice differences can strongly influence the extent of apparent proton delocalization (*52*). Notably, this sensitivity to minor environmental differences suggests that LBHBs are not as highly stabilizing as sometimes proposed (*45*). The high degree of structural similarity between DJ-1 and YajL (*53*) provides an opportunity to probe the influence of protein environmental effects on a Glu-Asp H-bond located in largely conserved but non-identical protein microenvironments in these homologs.

Our examination of the environment of the Glu-Asp pair identified two divergent residues between YajL and DJ-1 that might influence the degree of proton localization in this H-bond: Met17(DJ-1)/Thr16(YajL) and Ile21(DJ-1)/Thr20(YajL) (Figure 5A). In YajL, the Thr16 hydroxyl group donates an H-bond to the Oδ2 atom of Asp23, while this residue is Met17 in DJ-1. Also, YajL features an extended H-bonding interaction starting with Thr20, whose sidechain hydroxyl donates an H-bond to the peptide carbonyl of Thr16, which in turn donates a peptide amide H atom in an H-bond to Oε1 of Glu14 (Figure 5A). In DJ-1, these are non-polar residues that cannot H-bond in the same way, potentially providing a candidate mechanism by which sequence variation at these positions could alter the degree of proton delocalization in the Glu-Asp H-bond. Therefore, we individually introduced I21T and M17T substitutions in DJ-1 and determined each mutant’s effect on proton sharing at the Glu-Asp H-bond. X-ray bond length analysis of atomic resolution X-ray crystal structures indicates that M17T DJ-1 (1.00 Å resolution) retained a conventional H-bond at Glu15-Asp23, while I21T DJ-1 (0.97 Å resolution) had greater LBHB character, reflected in bond lengths that indicated both interacting oxygen atoms were protonated (Table 1).We note that M17T DJ-1 crystals were grown at pH=8.5, while I21T DJ-1 crystals were grown at pH=7.4, which could have some influence over H-bond protonation, although YajL crystals with an LBHB at the corresponding crystals are also grown at pH=8.5, indicating that full proton delocalization can occur in the highest pH used for DJ-1 crystallization and thus the presumed effect of pH differences in this range is minor. These mutations have moderate effects on DJ-1 thermal stability (Figure S2) that do not correlate clearly with degree of proton delocalization, likely because these non-conservative mutations alter several interactions that contribute to DJ-1 stability.

**Figure 5:**
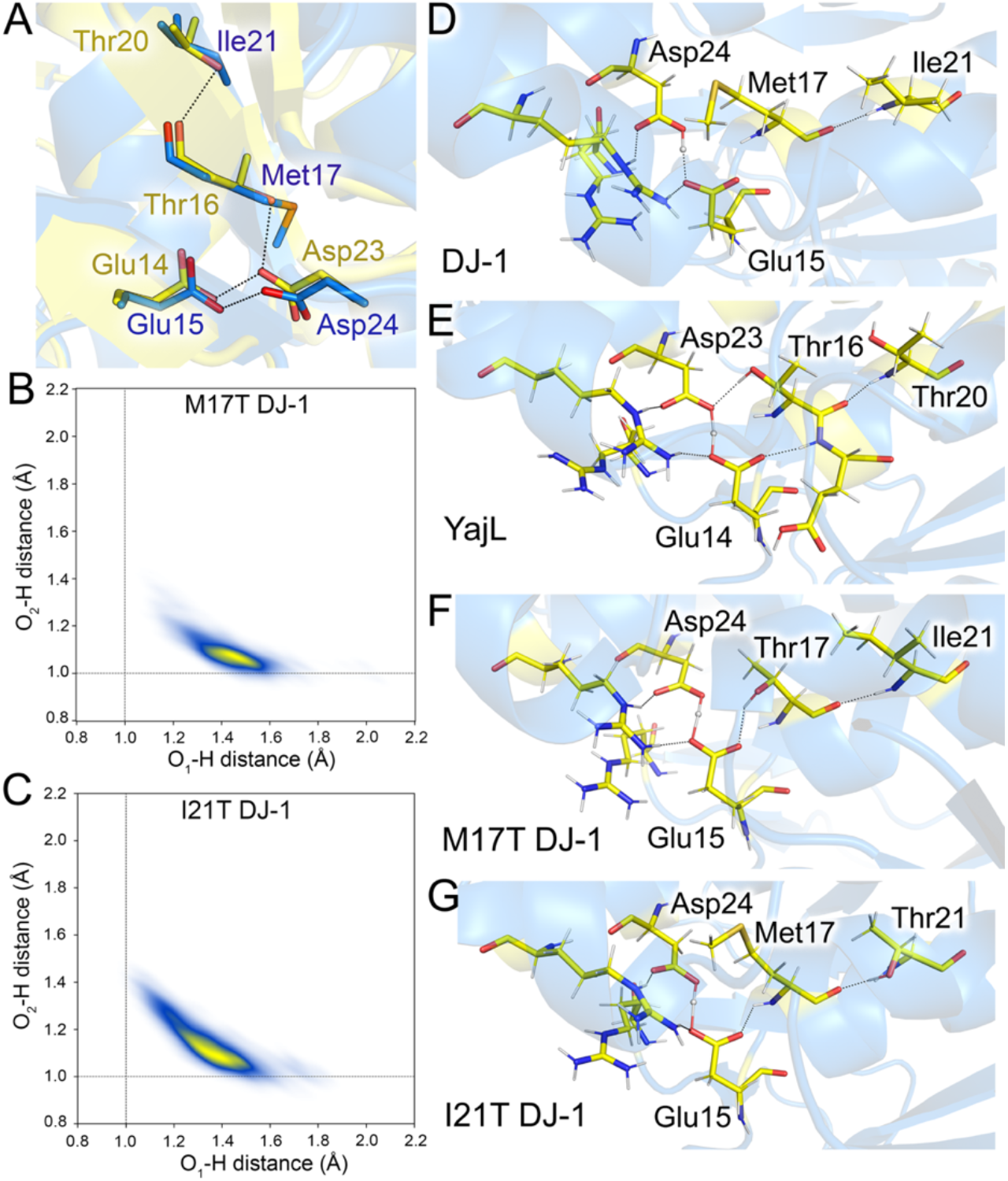
Microenvironmental differences between DJ-1 and YajL influence Glu-Asp proton localization. (A) Carton of the superposition of YajL (blue) and DJ-1 (yellow) show residues that differ in the extended H-bonding network that includes the Glu-Asp H-bond. YajL has more polar residues (Thr) at several locations. Panels B and C show QM/MM-BOMD simulation-derived density of states for the position of the proton in the Glu-Asp H-bond in two site-driected mutants of DJ-1 that make it more YajL-like. The I21T mutation results in partial delocalization of the proton (C), while the M17T mutation does not (B). Panels D-G show snapshots from the QM/MM-BOMD simulation of the wild-type DJ-1 and YajL as well as the two mutants of DJ-1. A common feature in YajL and I21T DJ-1 is the Glu residue accepting an H-hond donated by a peptide amide, correlating with greater proton delocalization in the Glu-Asp H-bond.

### Large QM/MM simulations capture environmental influence on proton localization in DJ-1 and YajL

For the engineered M17T and I21T DJ-1 mutants described above, we used quantum mechanics-molecular mechanics Born-Oppenheimer molecular dynamics (QM/MM-BOMD) to sample the equilibrium state of the Glu-Asp H-bond. QM/MM-BOMD simulations show that the degree of proton localization in this H-bond in M17T DJ-1 is similar to wild-type DJ-1, with an Asp24 donor O-H distance of ∼1.0 Å and a Glu15 acceptor distance of ∼1.4 Å (Figure 5B). By contrast, I21T DJ-1 has shifted its population to be more LBHB-like, with an increase in the donor distance to ∼1.2 Å, acceptor distance to ∼1.3 Å, and an overall broadening of proton locations over the course of the trajectory (Figure 5C). YajL still has a more delocalized proton, with a high population of protons ∼1.1-1.5 Å between both H-bonding residues, compared to I21T DJ-1 which samples more unevenly, localizing from ∼1.0-1.4 and ∼1.2-1.3 Å. This is consistent with the atomic resolution X-ray bond length analysis for both mutants (above).

For the M17T and I21T simulations, two different QM regions were used: a smaller one including only the Glu/Asp amino acids and a larger one that encompassed the central Glu-Asp H-bond as well as nearby residues including residues 17, 21, two waters, Arg28, and Arg48, which lie within ∼6Å of the Glu-Asp H-bond (see Methods). Within this larger region there are several conserved H-bonding interactions with the Glu-Asp pair in both YajL and DJ-1, including a salt bridge between Arg28 (DJ-1 numbering) and both the Oε2 atom of Glu15 (H-bond acceptor) and the Oδ1 atom of Asp24 (not involved in the H-bond) (Figure 5D,E). Also, in both DJ-1 and YajL, the active site Glu18 residue (DJ-1 numbering) backbone amide donates an H-bond to Glu15 Oε1, connecting this dimer-spanning Glu-Asp H-bond to the active site. A common feature in YajL and I21T DJ-1 that is not observed in DJ-1 and the M17T mutant is that the Glu14/15 residue in YajL and I21T DJ-1 accepts an H-bond donated by a nearby peptide amide, which correlates to greater proton delocalization in the Glu-Asp H-bond (Figure D-G). Although H-acceptor atom distances are similar between the small and large QM region simulations, the H-donor atom distances are highly influenced by a QM description of the surrounding environment (Figure S3). Therefore, an accurate quantum mechanical description of the local environment of the LBHB is crucial to understanding its influence on the localization of the proton in LBHBs. This is particularly true for distinguishing between more subtle environmental effects on proton localization observed for the M17T and I21T DJ-1 mutations (Figure 5F,G), where a larger QM region that included the nearby Arg dyad and intervening residues was needed for agreement with experiment, in contrast to the simulations of the wild-type protein where only the immediate environment of the Asp-Glu H-bond required QM treatment (Figure S3). In addition, we ran additional longer timescale simulations with a deuterium between the key Glu/Asp residues in the QM region (see Methods). The overall sharing of the deuteron in the Asp-Glu H-bond is essentially identical to hydrogen (Figure S4), with the period of vibration slowing by about one half, as expected for the increase in mass of the deuteron.

To obtain specific barrier heights for proton transfer from the Asp to Glu residue, we performed umbrella sampling calculations (Figure S5). These calculations provide a view of proton localization similar to the QM/MM-BOMD simulation, while also providing energies. In each case, we plotted the potential of mean force (PMF) as the proton is shifted from one residue (−0.4 to −0.2 Å from perfectly shared) to the other (+0.4 to +0.2 Å from perfectly shared), where perfectly shared refers to a proton precisely in between the donor and acceptor atoms (0 Å) (Figure 6A). Wild-type DJ-1 has an uphill barrier from donor to acceptor, indicating a fully localized proton on Asp24 and a barrier of over 6 kcal/mol to proton transfer to Glu15. M17T DJ-1 also has a high barrier to proton transfer between these residues (Figure 6A). By contrast, YajL has a nearly flat free energy profile with a small barrier to transfer of ∼1.1 kcal/mol and a local minimum almost exactly at 0.0 where the proton is perfectly shared, as expected for an LBHB. I21T DJ-1 appears to have mixed character—the proton would finish its transfer from Asp24 to Glu15 at +0.2 Å where the energy increases, indicating that the proton is still somewhat biased towards the donor Asp residue. However, the barrier to reaching the shared region of the reaction coordinate (∼0.0) in I21T DJ-1 is 1.15 kcal/mol, similar to the barrier in YajL. Therefore, in both types of simulation, YajL provides a clear example of a LBHB and shares potential energy surface features with I21T DJ-1 that indicate partial proton delocalization in that mutant.

**Figure 6:**
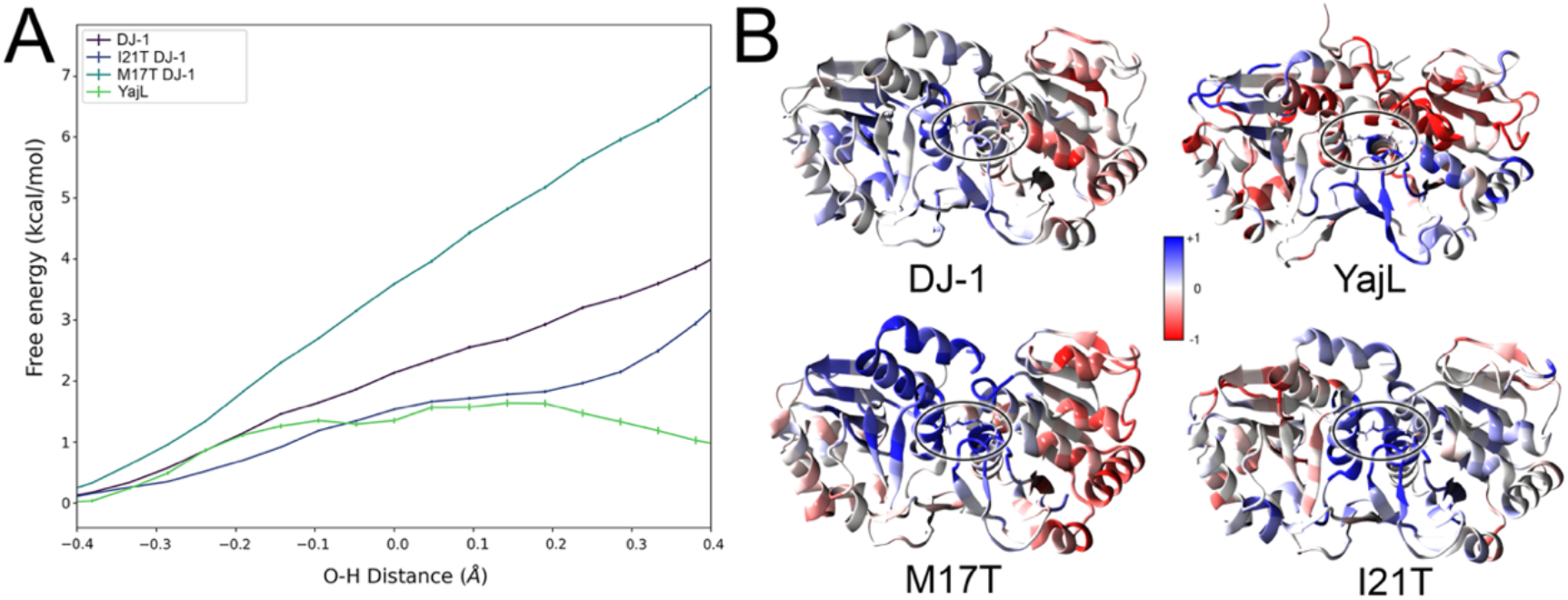
Energy barrier to proton transfer in the Glu-Asp H-bond and effects of proton sharing on correlated protein motions. Panel A shows the potential of mean force (PMF, Y-axis) for transfer of the proton from the Asp residue to the Glu residue in the indicated proteins. YajL (green) has a low barrier of ∼1 kcal/mol and similar energies for protonated Asp and Glu residues, I21T DJ-1 (dark blue) also has a low barrier but the proton is favored on the donor, while both DJ-1 (purple) and M17T DJ-1 (light blue) have higher barriers and strongly favored Asp protonation. Panel B shows the correlation of atomic motions in the QM/MM-BOMD simulation with proton motion in the Glu-Asp H-bond mapped onto the ribbon diagram of the dimers of each protein. The H-bond is circled. The degree of proton delocalization influences the extent and nature of dimer-spanning correlated motions. Correlation is mapped onto the protein ribbon diagrams and are colored blue (+1) for perfect correlation and red (−1) for perfect anti-correlation), as shown on the bar.

In addition to the influence of the local environment on the dimer-spanning Glu-Asp H-bond, QM/MM-BOMD simulations revealed an unexpected large-scale correlation of dimer motion relative to the proton transfer coordinate of the Glu-Asp H-bond. We extracted the cross-correlations of each protein atom relative to the proton transfer coordinate for each system during the simulation and mapped these correlation vectors onto the proteins (Figure 6B, S5). Wild type DJ-1 and M17T DJ-1, both of which have a localized proton in a conventional Glu-Asp H-bond, have similar positive and negative correlations across the dimer interface. In these two cases, the protomer associated with the hydrogen bond donor moves with the proton and the monomer associated with the acceptor moves against it (Figure 6B). By contrast, YajL and I21T DJ-1 exhibit a more LBHB-like Glu-Asp interaction and display a strong correlation of motion that spans the dimer interface and extends deeply into each monomer (Figure 6B). Specific secondary structures in each protomer, specifically α-helices that are adjacent to the dimer interface, move with the proton, and α-helices adjacent to the β-sheet region move opposite to the proton. Rather than independent motion, where each protomer is moving essentially as an independent unit, the correlation is coupled throughout the finer structural details of each protomer (Figure S6). This suggests that the proton movement in the LBHB is a mechanism that correlates motions in both monomers of the dimer, indicating that the LBHBs can tune the structural and dynamic aspects of the entire dimer interface.

## Discussion

Analysis of bond lengths in atomic resolution X-ray crystal structures has long been used to determine the protonation state of carboxylic acids (*17, 26, 29*). Because the lengths C-O bonds of protonated carboxylic acids differ by ∼0.1 Å (1.31 Å for C-OH vs 1.21 Å for C=O) while there are no differences in C-O bond lengths of ionized carboxylates (1.26 Å), carboxylic acid protonation states can be confidently determined once the estimated standard uncertainties on C-O bond lengths drop below ∼ 0.03 Å, i.e. above the 3σ threshold. This typically occurs once the resolution of the X-ray diffraction data extends beyond ∼1.2 Å (*29*). The application of bond length analysis to identify LBHBs between carboxylic acids was previously proposed by us (*18*) but has not been previously corroborated using other experimental techniques. Because both DJ-1 and YajL form large and well-diffracting crystals, they provide an opportunity to use neutron crystallography to validate atomic resolution X-ray bond length analysis as a tool for LBHB detection. Here, we found that both neutron diffraction and QM/MM-BOMD simulations corroborate X-ray bond length analysis, identifying the LBHB in YajL and the corresponding conventional short H-bond in DJ-1. Because COOH-^-^OOC hydrogen bonds have high potential to be LBHBs and atomic resolution macromolecular X-ray diffraction data are increasingly available, we echo our previous call that X-ray bond length analysis should be more widely applied to identify LBHBs in other systems if high-resolution X-ray data are available (*18*). As of November 2025, there are 7,594 X-ray crystallographic structures in the PDB with resolutions of 1.2 Å or better, providing a large dataset for exploration. In addition, recent advances in using quantum mechanics in the refinement target function have been used to identify the LBHB in YajL even when data are truncated to 2.0 Å (*30*), marking a major advance in the toolkit that can be used to identify LBHBs (and other unusual quantum mechanical phenomena) in X-ray diffraction datasets of average resolution in the PDB. We note that truncated data often provide greater signal than datasets that reach the same resolution using conventional criteria for data cutoff, and therefore the minimum resolution needed to extract this information using quantum refinement will need further exploration.

Perdeuteration of YajL and DJ-1 had negligible effect on the Glu-Asp H-bond using a statistically rigorous bond length analysis comparison, consistent with other studies showing generally small effects of deuteration on protein crystal structures (*46, 47, 54*). However, deuteration is known to have a substantial effect on protein stability and dynamics (*43*), causing changes in H-bonding structure and energetics (*55, 56*) that stabilize and rigidify proteins (*57, 58*). This effect may be evident in the X-ray diffraction data presented here, where perdeuterated DJ-1 and YajL in D_2_O consistently diffracted to higher resolution than their protiated counterparts. The robustness of the YajL LBHB to perdeuteration is noteworthy because LBHBs have anomalous H/D exchange kinetics (*48*), reflecting the influence of H->D substitution on the zero-point energy of an H-bond. More surprisingly, deuteration did have a measurable effect on the protonation state of Glu18 (DJ-1 numbering) in the active site of both enzymes. Glu18 is protonated at physiological pH in H_2_O and donates its H atom in an H-bond to the Cys106 thiolate S atom, contributing to the low pK_a_ value of the Cys106 thiol(ate) (*50*). The differential response of two classes of H-bond (COOH-^-^OOC vs COOH-^-^S) to deuteration suggests that the reduced donor atom-D zero-point energy can have variable energetic effects in distinct protein microenvironments and should be considered when collecting or interpreting biophysical data that use H/D substitution.

The requirement for matched donor and acceptor pK_a_ values in an LBHB suggests that proton delocalization in these bonds should be sensitive to nearby mutations, because pK_a_ values are highly influenced by the microenvironment of ionizable groups (*59*). Our results with DJ-1 mutants (particularly I21T) support a model where even distal residues participating in H-bond networks can significantly alter the barrier height to proton transfer in a carboxylic acid-carboxylate H-bond, converting a conventional short H-bond to a more LBHB-like interaction. QM/MM simulations with large QM regions were needed to capture these extended environmental effects in I21T DJ-1, allowing the proton behavior in the bond to evolve naturally during simulation without applying biasing or other artificial potentials. The ability of QM methods to reproduce extended environmental effects on proton delocalization indicates that future efforts could use large QM region simulations as an early step to evaluate candidate mutations *in silico* and then follow with experimental characterization. Such simulation capability is an essential step in developing the ability to engineer LBHBs, as it provides a predictive computational tool that can reduce time-consuming and expensive experimental screening of many candidate mutations.

The role of the LBHB in YajL is unknown. This Asp/Glu interaction is conserved from bacteria to humans and spans the dimer interface, suggesting an important role in either maintaining structure or function of these DJ-1 family proteins. A conservative Asp->Asn mutation in YajL decreases the protein thermal stability of by 4.0° C, which is significant. Moreover, the difference in degree of proton delocalization between YajL and DJ-1, which are close structural homologs despite billions of years of evolutionary divergence, suggest either that this H-bond has divergent functions in these proteins or that the degree of proton delocalization in the resting state of these enzymes is variable or of secondary importance. The Glu-Asp pair is ∼6Å from Glu18, an important residue in the active site of DJ-1 family proteins that is proposed to be a general acid/base in the glyoxalase (*60*) and cyclic phosphoglycerate anhydride hydrolase activities (*61*). Speculatively, the proximity of the Glu-Asp H-bonded pair to another carboxylic acid with perturbed pK_a_ in the active site suggests a mechanism by which this unusual H-bond could be indirectly connected to catalysis. Furthermore, the correlation between the degree of proton delocalization and the dynamics of the protein dimers was unexpected and suggests that LBHBs could exert a subtle but potentially important influence on long-range correlated protein dynamics, including communication across the dimer interface. This dynamic effect adds to the known importance of LBHBs in facilitating proton mobility in multimeric enzymes (*8*) and photosystem II (*62*), establishing the ability of LBHBs to allosterically influence protein function. Our results indicate that the extent of proton (de)localization exists along a continuum in different carboxylic acid-carboxylate H-bonds and is acutely sensitive to environmental influences. We propose that this environmental sensitivity makes carboxylic acid-carboxylate H-bonds attractive targets for engineering charge-assisted H-bonds or LBHBs in proteins that facilitate proton transfer reactions.

## Materials and Methods

### Protein expression and purification

The full-length sequences of *Escherichia coli* YajL and human DJ-1 were cloned between the NdeI and XhoI sites of the bacterial expression vector pET15b (*53, 63*) and expressed in BL21(DE3) Escherichia coli. Mutations I21T and M17T in DJ-1 and D23N in YajL were generated by site-directed mutagenesis PCR. To produce hydrogenated proteins, bacteria were grown in LB medium supplemented with 100 μg/ml ampicillin (Fisher Scientific) at 37 °C with shaking. Protein expression was induced by adding 0.5 mM isopropyl β-d-1-thiogalactopyranoside (IPTG) (Goldbio) once the culture A_600_ reached 0.6-0.8. After 4 hours, the induced culture was harvested by centrifugation, and the cell pellets were stored at −80 °C until needed.

To produce perdeuterated proteins, a frozen glycerol stock was first revived in H_2_O minimal medium with unlabeled glycerol (0.5% w/v) and carbenicillin (100 μg/mL) because frozen stock cultures do not passage well directly into deuterated media (*64*). After initial growth in H_2_O minimal medium, the cells were adapted stepwise to minimal medium containing increasing percentages (50,75, and 100%) of D_2_O. Unlabeled glycerol (0.5% w/v) was used as the carbon source in all media except the 100% D_2_O minimal medium, which used glycerol-d_8_ (0.5% w/v). The final D_2_O-adapted preculture was used to inoculate the bioreactor vessel to an initial volume of 1.4 L; final carbenicillin concentration was 100 μg/mL. Following inoculation, the BioFlo 310 bioreactor controller console (Eppendorf, Enfield, CT) was set to 30 °C and the dissolved oxygen to >30%. The pD was kept above 7.3 by the controlled addition of sodium deuteroxide solution in D_2_O (10% w/w). Once the initial glycerol-d_8_ was exhausted, the culture was fed with a solution containing 10% (w/v) glycerol-d_8_, 0.2% (w/v) MgSO_4_, and carbenicillin (100 μg/mL). Protein expression was induced at an OD_600_ of ∼9 by addition of isopropyl β-d-1-thiogalactopyranoside (IPTG) to a final concentration of 1 mM. The cells were collected ∼14 h later by centrifugation at 6,000 x g for 40 min. After removing the supernatant, the wet cell paste was harvested and stored at −80 °C until further use.

Pelleted cells were resuspended in a solution containing 50 mM HEPES pH 7.5, 300 mM NaCl, 2 mM DTT, and 1mg/ml lysozyme, and then lysed by sonication. The resulting lysates were clarified and applied to a His-Select Nickel affinity resin (Sigma) for protein purification. Recombinant proteins tagged with a His_6_ tag were eluted from the resin using 250 mM imidazole. The N-terminal hexahistidine tag was removed by thrombin cleavage at 4 °C for 12 hours, followed by sequential passage over His-Select resin to remove any remaining tagged protein and benzamidine-Sepharose resin to remove the thrombin. All purification solutions were prepared using H_2_O.

For protiated proteins, the purified protein was concentrated to 22mg/ml in storage buffer (25 mM HEPES, pH 7.5, 100 mM KCl, 2.5 mM DTT) before being snap-frozen on liquid nitrogen and stored at −80 °C. For perdeuterated proteins, solvent was exchanged by dilution with at least ten volumes of D_2_O storage buffer, incubated for 12 hours at 4 °C, and then concentrated. This process was repeated six times, resulting in an exchange factor of ∼10^6^. The proteins were concentrated to 22mg/ml in D_2_O storage buffer, snap-frozen on liquid nitrogen, and stored at −80 °C.

### Thermal stability assay

The thermal stabilities of wild-type and D23N YajL were determined using the Thermofluor assay (*65*). Protein samples at concentrations ranging from 1-5 mg/mL were mixed with Sypro Orange dye (Sigma-Aldrich) and placed in optically clear, thin-walled PCR tubes (Eppendorf) The mixture was heated from 20 to 95°C at 2°C/min using an iCycler iQ real-time thermal cycler. Fluorescence was measured at an excitation wavelength of 490 nm and an emission wavelength of 575 nm. Fluorescence data were plotted as the first derivative with respect to temperature, and the peak of this curve indicates the reported melting temperature (T_m_). The assay was performed in triplicate at different protein concentrations, and representative data are shown in Figure 3.

The thermal stabilities of mutants of DJ-1 were determined by circular dichroism (CD) at 222 nm. Protein samples were prepared at 0.4 mg/mL (20 μM monomer concentration) by dilution and dialysis against 10 mM sodium phosphate buffer (pH=7.4) and then placed in a 0.1 cm path length quartz cuvette. Measurements were carried out on a Jasco J-815 CD spectrometer with a Jasco CDF-426S/15 temperature control system. The temperature was increased from 20°C to 95°C at 5°C/min. The thermal denaturation curves were plotted as ellipticity versus temperature, and the reported melting temperatures (T_m_) were determined by fitting the data to a sigmoidal dose-response curve.

### Protein crystallization for X-ray and neutron diffraction

For X-ray crystallography, DJ-1 and YajL proteins were crystallized using the sitting drop vapor diffusion method by mixing 2 μL of protein and 2 μL of reservoir solution and equilibrating against 500 μl of reservoir solution at room temperature (∼22°C). DJ-1 crystals in space group P3_1_21 typically formed overnight against a reservoir solution containing 22-24% PEG 3000, 200 mM NaCl, and 100 mM HEPES, pH 7.5 and were left to grow for approximately two weeks. M17T DJ-1 crystals were grown against a reservoir solution containing 19-22% PEG 4000, 200 mM MgCl_2_, and 100 mM Tris HCl, pH=8.5. I21T DJ-1 crystals were grown against a reservoir solution containing 22-27% PEG4000, 0.2 M sodium acetate, and 100 mM Tris HCl, pH=7.4. YajL crystals were grown against a reservoir solution containing 19-22% PEG 4000, 225 mM MgCl_2_, and 100 mM Tris HCl, pH 8.1. D23N YajL crystals were grown from a reservoir solution containing 22-25% PEG 4000, 200 mM MgCl_2_, and 100 mM Tris, pH=8.5. For YajL and D23N YajL crystals, microseeding using a cat whisker was used to improve crystal quality by reducing the size of the hollow depletion zones at the ends of the prismatic crystals. Perdeuterated DJ-1 crystals used for X-ray diffraction were grown in identical conditions to regular DJ-1 crystals, with the exception that all solutions were prepared using D_2_O. Perdeuterated YajL crystals were grown against a reservoir solution consisting of 16-18% PEG 4000, 225 mM MgCl_2_, and 100 mM Tris HCl, pH=8.1; all solutions were prepared using H_2_O as crystals did not grow well against D_2_O-containing buffers. As above, microseeding was used to enhance the quality of YajL crystals. Prior to harvesting the YajL crystals, H/D exchange was performed by replacing the reservoir buffer with D_2_O-containig buffers and allowing for vapor equilibration *in situ* for 4 or more weeks.

For neutron diffraction, perdeuterated DJ-1 was crystallized in a 9-well glass plate and sandwich box set-up with 200 µL drops of protein mixed with 20% PEG3350, 100 mM HEPES pH=7.5 and 200 mM KCl at 1:1 ratio. After ∼1 month of incubation at 18 °C, the temperature was reduced to 16 °C, and the crystals continued to grow for two more months. Long, hexagonal javelin-shaped crystals of perdeuterated DJ-1 in space group P6_5_22 grew in solutions made with H_2_O, whereas bipyramidal crystals in space group P3_1_21 grew in D_2_O solutions. DJ-1 crystals in space group P6_5_22 were mounted in a fused quartz capillary accompanied with 22% PEG 3350 prepared with 100% D_2_O and allowed to H/D exchange for several weeks before beginning neutron diffraction data collection. Crystallization of perdeuterated YajL (20 mg/mL) was achieved similarly, however crystals grew better in Hampton microbridges using 100 mM Tris pH=8.0, 250 mM MgCl_2_ and 20% PEG 4000 made with H_2_O as the well solution. H/D exchange was first performed *in situ* by replacing the well solution with the one made with D_2_O and keeping a crystal in the crystallization drop for two weeks. After H/D exchange, the crystal was mounted in a fused quartz capillary accompanied with 22% PEG4000 prepared with 100% D_2_O for neutron diffraction.

### Neutron and X-ray diffraction data collection, processing and structure refinement

For X-ray diffraction, crystals were cryoprotected in their respective reservoir conditions supplemented with an additional 25-30% ethylene glycol and then cryocooled in liquid nitrogen as previously described (*18*). Synchrotron X-ray data collection was performed at beamline 12-2 at the Stanford Synchrotron Radiation Lightsource (SSRL) using shutterless data collection, 0.729 Å (0.886 Å for YajL D23N) wavelength incident X-rays, and a Dectris Pilatus 6M pixel-array detector. To minimize experimental differences between datasets, all X-ray diffraction datasets for protiated and perdeuterated DJ-1 and YajL were collected during the same beamtime at SSRL beamline 12-2 with minimal changes in data collection configurations between samples. Diffraction data were collected at 100 K, indexed and integrated using XDS (*66*) and scaled using Aimless (*67*). Data statistics are presented in Table S1.

Neutron diffraction data were collected from perdeuterated DJ-1 and YajL crystals after preliminary screening for diffraction quality using a broad-bandpass Laue configuration using neutrons from 2.8 to 10 Å at the IMAGINE instrument at the High Flux Isotope Reactor (HFIR) at Oak Ridge National Laboratory (*68-70*). The full neutron diffraction datasets were collected using the Macromolecular Neutron Diffractometer (MaNDi) instrument at the Spallation Neutron Source (SNS) (*71, 72*). The crystals were held stationary at room temperature and diffraction data were collected for 20 hours using neutrons between 2.00-4.16 Å wavelength. During each experiment, the next 20-hour diffraction image was collected after crystal rotation by Δϕ = 10°. MaNDi neutron diffraction data were reduced using the Mantid package, with integration carried out using three-dimensional TOF profile fitting (*73*). Wavelength normalization of the Laue data was performed using the Lauenorm program from the Lauegen suite (*74-76*). The neutron data collection statistics are shown in Table S1.

Following neutron data collection, room-temperature X-ray diffraction datasets were collected from the same perdeuterated crystals at the same temperature used for neutron data collection. YajL X-ray data were collected on a Rigaku HighFlux HomeLab instrument equipped with a MicroMax-007 HF X-ray generator, Osmic VariMax optics, and an Eiger R 4M hybrid photon counting detector. DJ-1 X-ray diffraction data were collected on a Rigaku MicroMax-007 X-ray generator, Osmic confocal optics, and an RaxisIV^++^ image plate detector. The YajL diffraction data were integrated using the CrysAlis Pro software suite (Rigaku Inc., The Woodlands, TX) and the DJ-1 data were indexed using MOSFLM (*77*). Both datasets were scaled using Aimless (*67*) in the CCP4 suite (*78*). The room-temperature X-ray data statistics are shown in Table S1. Joint X-ray/neutron refinement was performed using PHENIX 1.19.2 (*79, 80*), with the X-ray structure serving as the starting model. All deuterium atoms were automatically added to the model using ReadySet in PHENIX and H/D occupancies were refined for exchangeable H/D atoms. Refining H/D atoms using the riding hydrogen model produced less overfitting and a smaller R_free_-R gap than did the individual sites model, consistent with expectation and prior results (*81*). However, we used a final individual H/D atom joint refinement to obtain occupancies and B-factors for the shared deuterons in the Glu-Asp H-bond, as they do not belong to a residue and therefore cannot be fully refined using the riding hydrogen model. We note that a newer implementation of joint refinement was released in PHENIX (*81*) which may address this issue. Cys106 in DJ-1 was modeled as partially oxidized to the Cys-SO_2_^-^, consistent with prior results (*82*). The final models were validated using the MolProbity server (*83*) and the PDB validation tools and model statistics are reported in Table S2.

Initial phases for DJ-1 and YajL were determined using molecular replacement with Phaser as implemented in CCP4 with PDB codes 1P5F (*84*) and 2AB0 (*53*), respectively. The model coordinates and anisotropic atomic displacement parameters (ADPs) were refined with stereochemical and anisotropic ADP restraints using Refmac5 (*85*) and phenix.refine (*79*). After each refinement cycle, the protein and solvent models were manually improved in Coot (*86*) based on the 2mF_o_ – DF_c_ and mF_o_ − DF_c_ electron density maps. We note the presence of electron density near Arg48 in DJ-1 that was conservatively modeled as water but has features that suggest it is another unknown species, possibly anionic. In addition, the I21T DJ-1 X-ray structure is modeled with Cys106 partially oxidized to Cys-SO_2-_and there difference electron density around Cys106 in both YajL and DJ-1 structures indicating photo oxidization, which has been noted to occur in DJ-1 superfamily proteins previously (*87*). The identities of modeled chloride ions in YajL were confirmed using anomalous difference electron density maps. The refined H-DJ-1 and H-YajL models were used as a starting point for rigid body refinement for the D-DJ-1 and D-YajL structures. The final models were validated using the MolProbity server (*83*) and the PDB validation tools and model statistics are reported in Table S2. As is typical of DJ-1/YajL structures, residue Cys106 is a Ramachandran outlier.

### Atomic resolution X-ray bond length analysis

Unrestrained bond lengths and their associated ESUs were determined by refining converged structural models in SHELXL (*88*) as previous described (*18*). Briefly, all geometric restraints (SHELX instructions DFIX, DANG) were removed from Asp or Glu residues and the model was refined using conjugate gradient least squares (SHELX instruction CGLS) for 20 cycles with geometric restraints for other residues and ADP restraints for all residues. The resulting model was subjected to a final cycle of fully unrestrained least-squared refinement (SHELX instruction L.S.) and ESUs on bond lengths were determined by inversion of the Hessian matrix with the BLOC1 instruction that includes only the coordinate degrees of freedom in the Hessian matrix. The sigma level of differences in the two C-O bond lengths in Asp/Glu were determined using the propagation of errors formula: Δ(C-O)/(σ_1_^2^+σ _2_^2^)^1/2^, where Δ(C-O) is the difference between the two C-O bond lengths in the carboxylic acid, σ_1_ is the ESU for the Cδ/γ-Oγ/ε1 bond and σ_2_ is the ESU for the Cδ/γ-Oγ/ε2 bond (*28, 29*).

### Protein molecular dynamics using QM/MM simulation

All four protein systems (DJ-1, YajL, M17T DJ-1 and I21T DJ-1) were prepared for simulation starting from their crystal structures by initial checking and protonation with MolProbity (*83*), pK_a_s of the side chains were calculated with propka3 (*89*) except for whose ionization states are experimentally known. Each system was solvated with a 12 Å pad of TIP3P water (*90*) to create a box size of ∼75 x 97 x 98 Å (approximately 7800 total waters) and neutralized with 4 Cl^-^ ions.

Classical MD was run with the pmemd.cuda module in Amber20 (*91, 92*), with a Langevin thermostat (*93*) in the NPT ensemble with a 5 ps^-1^ coupling constant and a 10 Å nonbonded interaction cutoff. Smooth particle mesh Ewald (*94*) was used to compute electrostatics with a 1 fs timestep, and hydrogen bond lengths were constrained using the SHAKE algorithm. We note that the classical force field description of the LBHBs is not stable without constraints, resulting in dimer dissociation after 50 ns. We selected snapshots prior to this point that resemble the crystal structure O-H-O distances in the dimer for QM/MM.

Born-Oppenheimer molecular dynamics with QM/MM (BOMD) was performed with the TeraChem 1.9.3 and OpenMM7 interface (*95-98*) . There were two QM regions chosen: one with a minimal QM region (only the Glu/Asp amino acids at one side of the dimer interface) and a larger QM region that included proximal H-bonding residues, mutation site, and key arginine residues (Arg28 and 48) that lie above the Glu/Asp residues (71 atoms total). We used the ωPBE hybrid functional (ω=0.22 Bohr-1) with a 6-31g** basis set. Each trajectory was run for at least 4 picoseconds with a 1 fs timestep. The force field parameters were the same as described above, except that the solvation was cut into a sphere (∼5000 waters) and spherical boundary conditions were applied with a restraining force of 10 kcal/mol * Å. We ran additional BOMD simulations with the same settings but deuterating the hydrogen at the key O-H-O distance, for 10 ps per system.

QM/MM umbrella sampling to obtain potential of mean force (PMF) for the proton transfer was performed with the TCPB interface between TeraChem 1.9.3 and Amber20. In this case, due to computational expense the basis set was lowered to 6-31g*, but the DFT functional was the same (ωPBE, ω=0.22). Harmonic restraints were applied for each window for the Glu-O—H—O-Asp distance, proceeding in 0.05 Å increment windows from 1.0 to 1.65 Å for 2.5 ps per window (23 windows totaling 57.5 ps per system, 230 ps of QM/MM sampling in total). The restraints were applied as a square potential with a value of 100 kcal/mol. The force field parameters remained the same. The Amber simulation parameters used were a Langevin thermostat with a 5.0 ps-1 collision frequency, no periodic boundary conditions, in the NVE ensemble and a Monte Carlo barostat, a 10 Å cutoff and a 0.5 fs timestep. The larger QM region (excluding backbone atoms) was used, including residues 15, 212, 48, 216, 16, 17 and the mutation point.

## Acknowledgments

We thank Jon Askey and Dr. Jennifer Clarke (U. Nebraska-Lincoln) for discussions and preliminary analysis of H-bonding statistics.

## Author contributions

J.L. purified protein, grew protein crystals, performed atomic resolution X-ray bond length analysis, and wrote the manuscript, O.G. and K.L.W. performed protein perdeuteration, D.W.K and A.K. grew protein crystals for neutron diffraction, L.C. and A.K. collected and processed room-temperature X-ray and neutron diffraction data and analyzed X-ray and neutron diffraction data, A.K. wrote the manuscript, M.A.H., S.Y.E., and A.R.W. performed computational work and analysis, and A.R.W. wrote the manuscript and obtained funding. M.A.W. designed the study, collected and analyzed X-ray and neutron diffraction data, wrote the manuscript, and obtained funding.

