## supplemental figures and tables for "Environmental Contributions to Proton Sharing in Protein Low-Barrier Hydrogen Bonds"

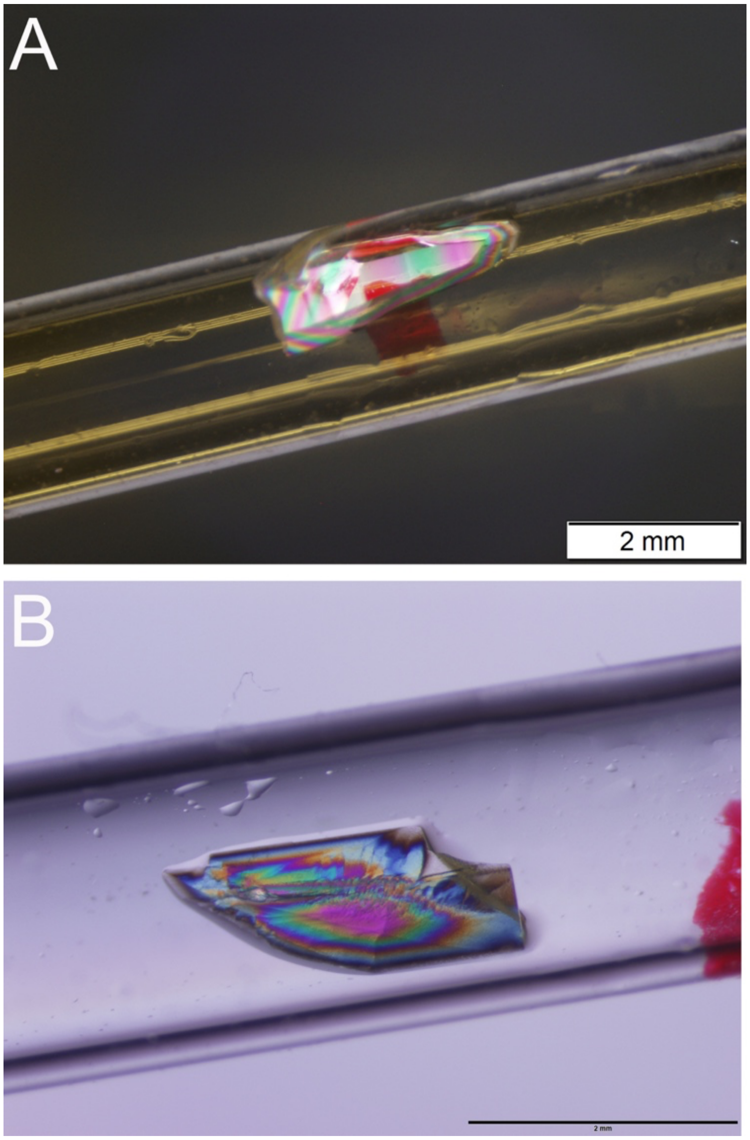


**Figure S1: Crystals of DJ-1 and YajL used for neutron diffraction.** Panel A shows a perdeuterated DJ-1 crystal in space group $P6_522$ with scale bar indicated. Panel B shows a perdeuterated YajL crystal in space group $P2_12_12_1$ with scale bar indicated. In both panels the samples shown were used to collect both neutron diffraction and accompanying room-temperature X-ray diffraction datasets, which were combined for subsequent neutron/X-ray joint refinement.

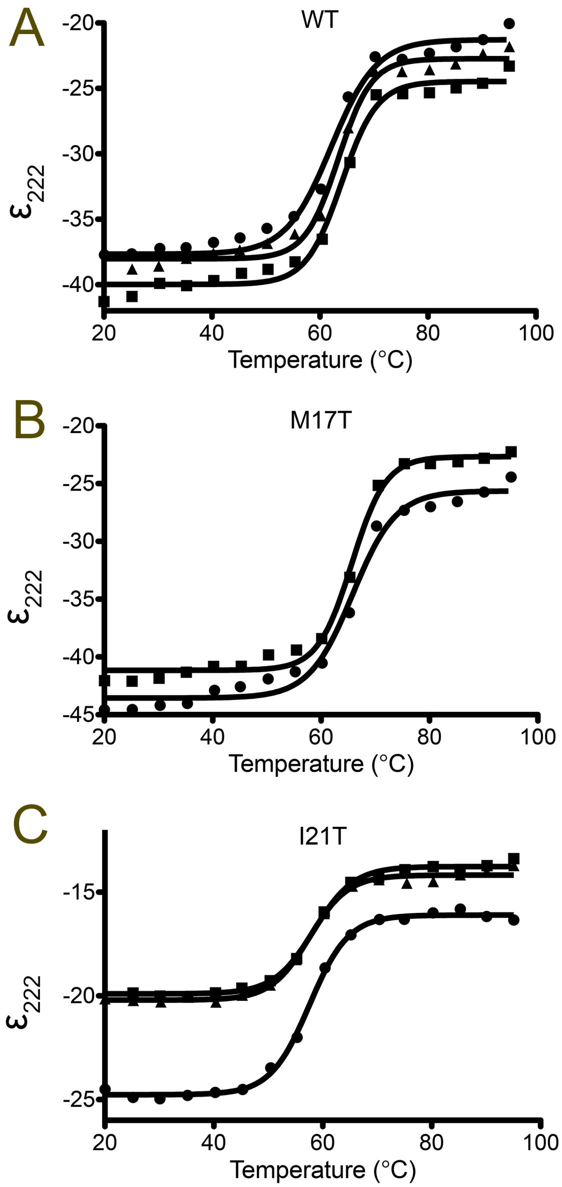


**Figure S2: Circular dichroism (CD) melting curves of DJ-1 mutants.** CD was used to monitor thermal melting of proteins using ellipticity at 222 nm ($\varepsilon_{222}$), with each replicate shown. Panel A shows wild-type DJ-1 measured in triplicate with average $T_m$(WT)=63.3°C. Panel B shows M17T DJ-1 thermal melting measured in duplicate with average $T_m$(M17)=65.7°C. Panel C shows I21T DJ-1 thermal melting curve measured in triplicate with average $T_m$(I21T)= 57.8°C.

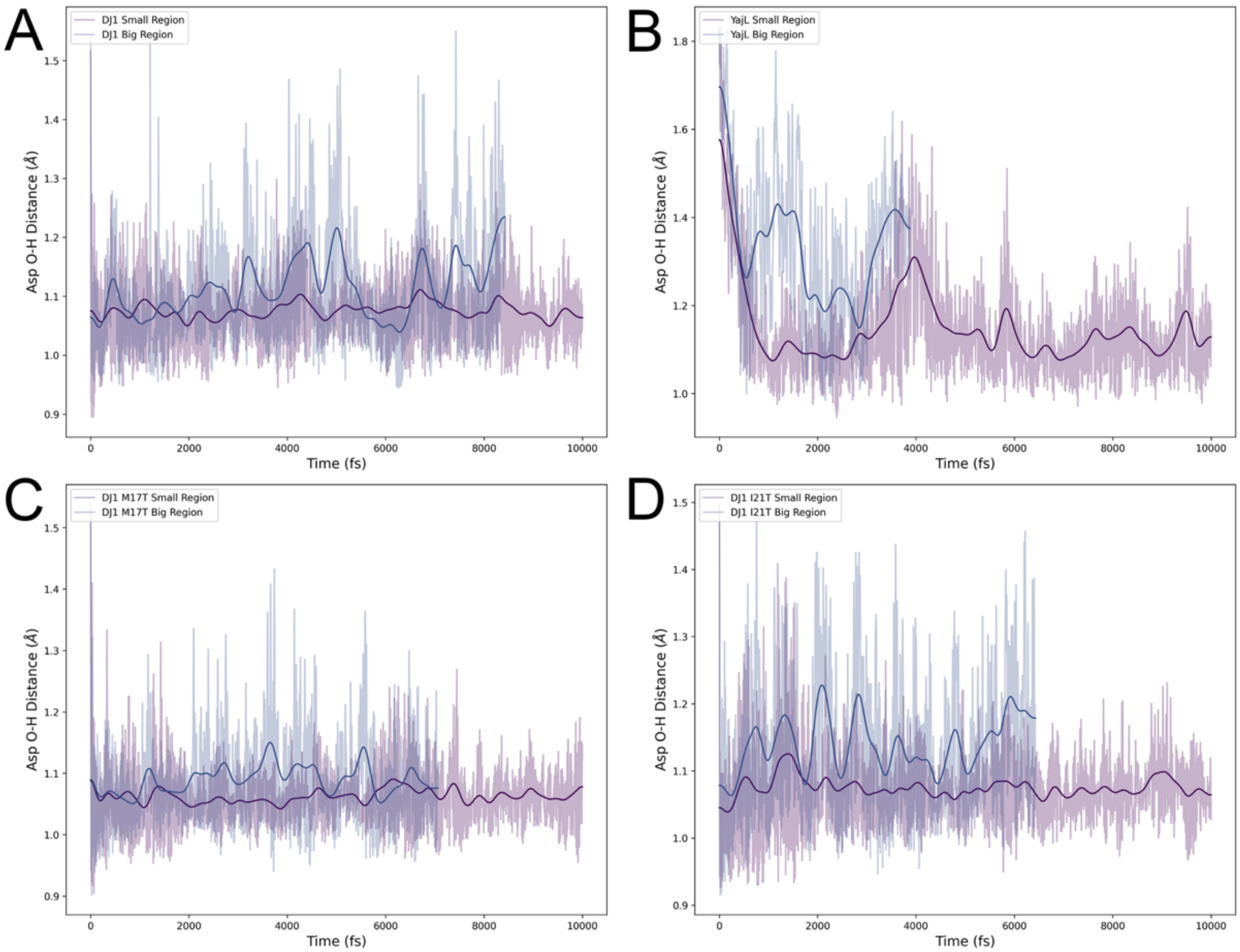


**Figure S3: Influence of smaller and larger QM regions on H-bond behavior in DJ-1, YajL, and DJ-1 mutants.** Key O-H distance of Asp24 plotted over time for YajL, DJ-1 and DJ-1 mutants with small QM region (purple) and large QM region (blue). The trajectories show large discrepancies particularly for LBHB-character systems.

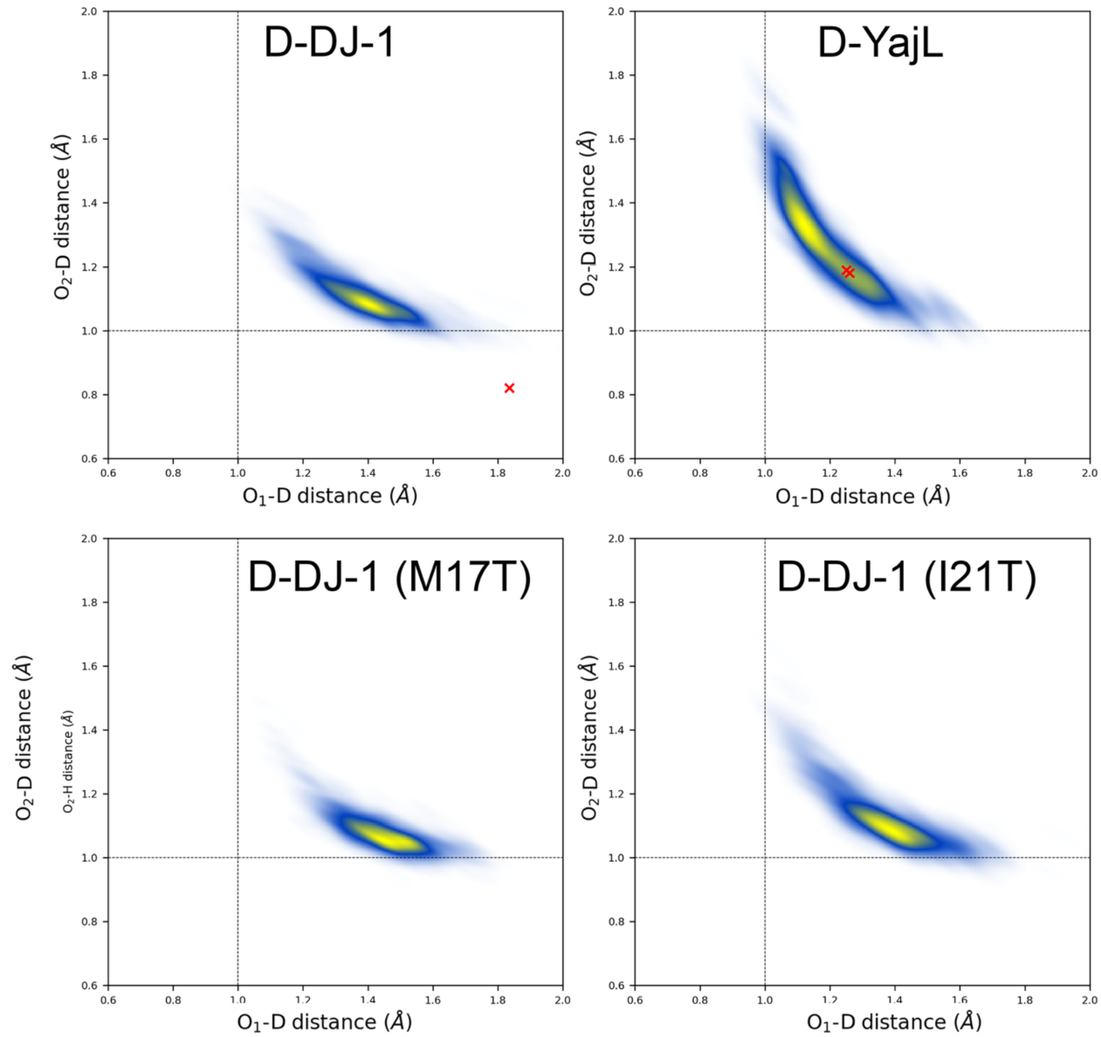


**Figure S4: Density of states for deuteron location in QM/MM-BOMD simulations of the dimer-spanning H-bond in deuterated proteins.** QM/MM-BOMD simulation-derived density of states for the position of the deuteron in the Glu-Asp H-bond in the indicated proteins, with the red crosses indicating the experimentally determined location of the deuterons from neutron diffraction. As with protiated protein simulations, the D-DJ-1, simulation shows a hydrogen atom localized predominantly on Asp24 (corresponding to the $O_2D$ distance on the Y-axis). By contrast YajL shows the deuteron localized more symmetrically between the oxygen atoms of Glu14 ($O_1$) and Asp23 ($O_2$), consistent with an LBHB. As with protiated proteins, I21T DJ-1 shows partial deuteron delocalization and M17T shows deuteron localization on Asp24. There are no significant differences between the behavior of the deuteron and the proton in this Glu-Asp bond.

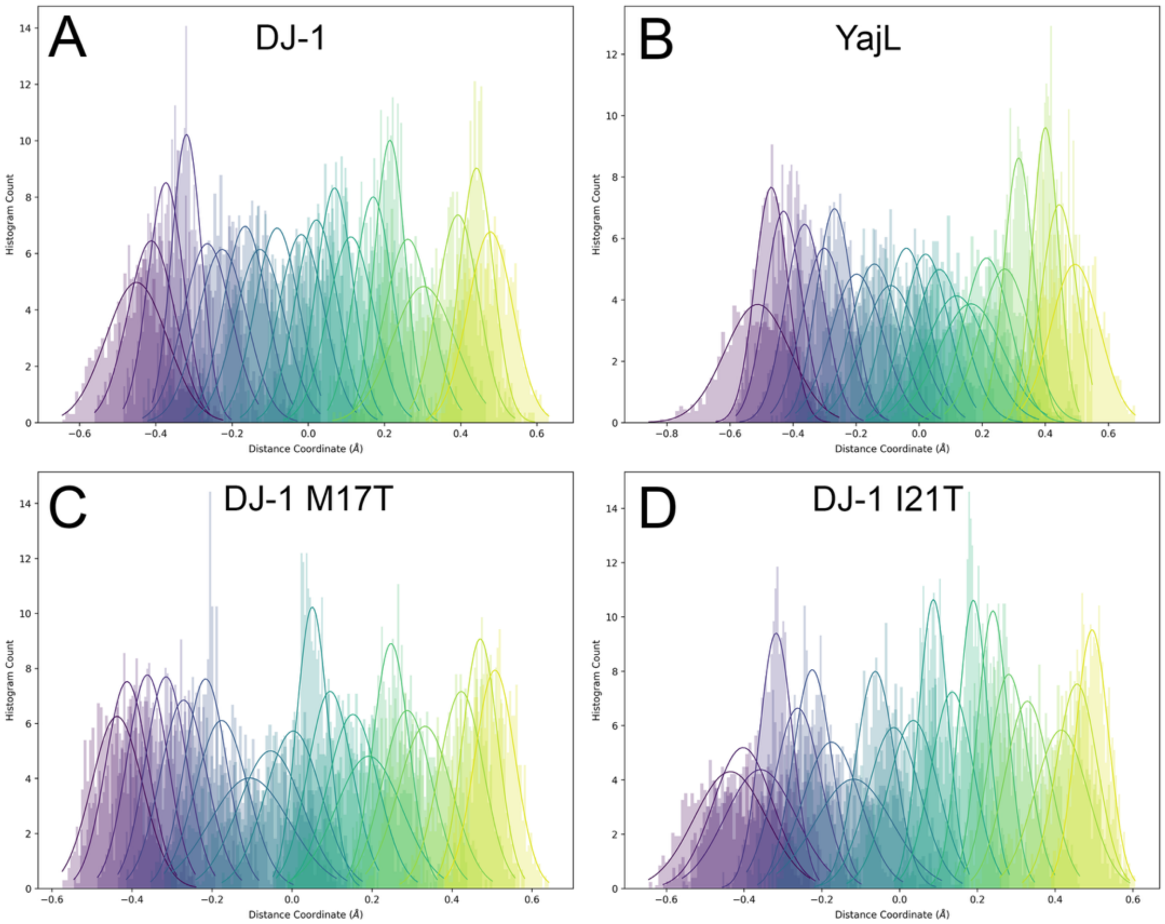


**Figure S5: Umbrella sampling histograms for DJ-1, YajL, and DJ-1 mutants.** Histograms of the O-H-O distances for Asp24-Glu14 for each umbrella sampling window for each system, colored from purple to yellow across the distance coordinate. Gaussian curves are fit to the maxima of each histogram. A distance of 0.0 corresponds to the center of the hydrogen bond (exactly 1.2 Å) between Asp24/Glu14. The histograms show sufficient overlap for adequate sampling of each distance.

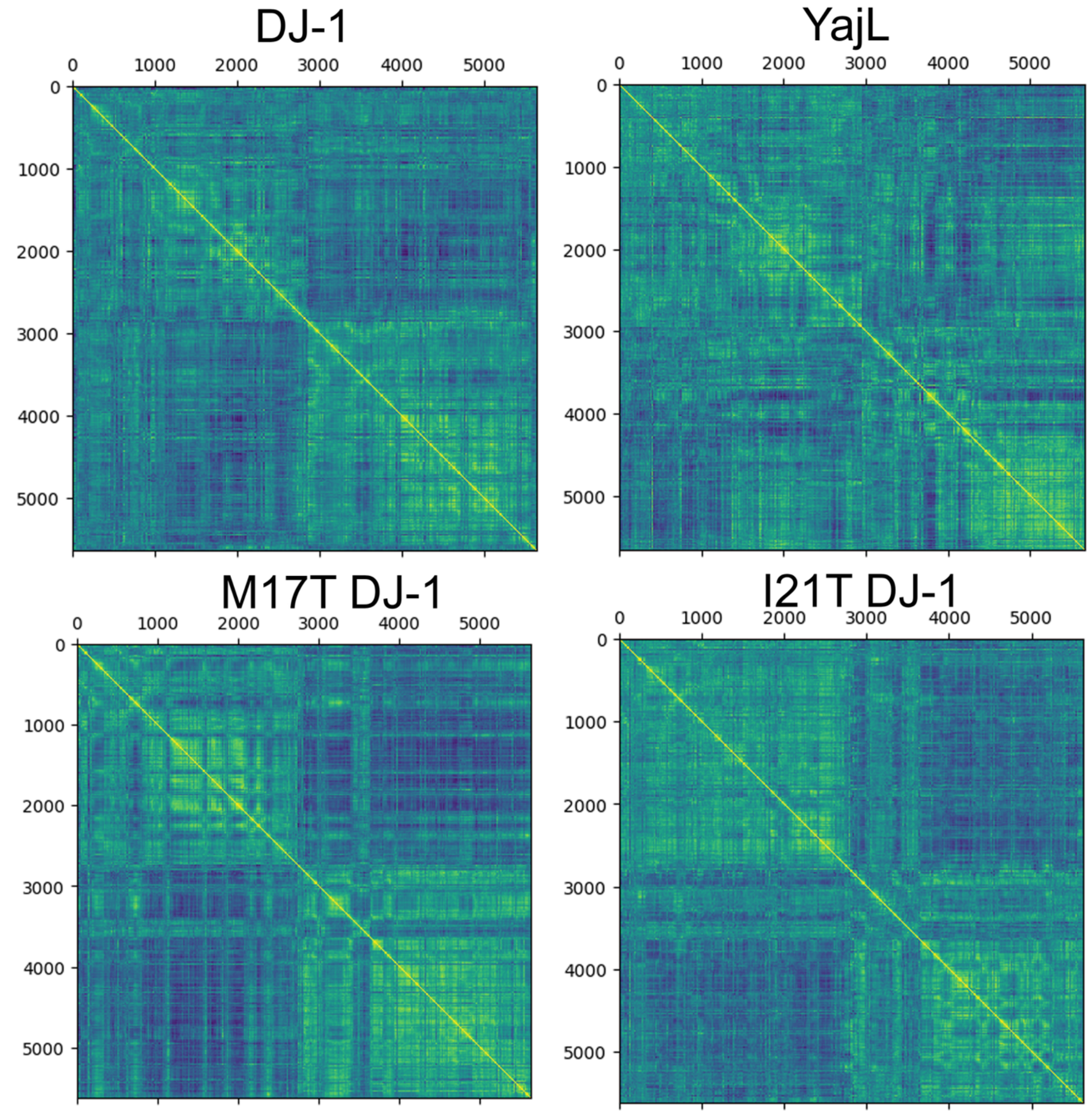


**Figure S6. Correlation plots for each protein atom against every other protein atom computed using the QM/MM BOMD trajectories**. Fully correlated atoms (normalized to 1.0) are shown in yellow, fully anticorrelated shown in blue (normalized to -1.0), and uncorrelated shown in green. The overall atomic correlation differs substantially between LBHB (YajL, I21T DJ-1) and normal H-bonded systems (wild-type and M17T DJ-1).

**Table S1:** Crystallographic data statistics

| Sample | H-DJ-1 | D-DJ-1 | H-YajL | D-YajL | DJ-1 M17T | DJ-1 I21T | YajL D23N | DJ-1 neutron | YajL neutron | D-DJ-1 RT | D-YajL RT |
|---|---|---|---|---|---|---|---|---|---|---|---|
| Diffraction source | SSRL 12-2 | SSRL 12-2 | SSRL 12-2 | SSRL 12-2 | SSRL 12-2 | SSRL 12-2 | SSRL 12-2 | ORNL MaNDI | ORNL MaNDI | Rigaku MicroMax-007 | Rigaku Highflux Homelab |
| Wavelength (Å) | 0.729 | 0.729 | 0.729 | 0.729 | 0.729 | 0.729 | 0.886 | 2.00-4.16 | 2.00-4.16 | 1.542 | 1.542 |
| Temperature (K) | 100 | 100 | 100 | 100 | 100 | 100 | 100 | 293 | 293 | 293 | 293 |
| Detector | Pilatus 6M pixel array detector | Pilatus 6M pixel array detector | Pilatus 6M pixel array detector | Pilatus 6M pixel array detector | Pilatus 6M pixel array detector | Pilatus 6M pixel array detector | Pilatus 6M pixel array detector | Spallation Neutron Source Anger camera detectors | Spallation Neutron Source Anger camera detectors | RaxisIV$^{++}$ | Eiger R 4M |
| Space group | $P3_121$ | $P3_121$ | $P2_12_12_1$ | $P2_12_12_1$ | $P3_121$ | $P3_121$ | $P2_12_12_1$ | $P6_522$ | $P2_12_12_1$ | $P6_522$ | $P2_12_12_1$ |
| a, b, c (Å) | 75.11<br>75.11<br>75.50 | 74.84<br>74.84<br>75.35 | 43.77<br>78.47<br>99.62 | 43.73<br>78.41<br>99.51 | 75.05<br>75.05<br>75.43 | 74.96<br>74.96<br>75.46 | 43.72<br>78.33<br>99.61 | 67.71<br>67,71<br>179.29 | 44.37<br>78.91<br>100.2 | 67.41<br>67.41<br>179.79 | 44.29<br>79.30<br>100.18 |
| α, β, γ (°) | 90.00<br>90.00<br>120.00 | 90.00<br>90.00<br>120.00 | 90.00<br>90.00<br>90.00 | 90.00<br>90.00<br>90.00 | 90.00<br>90.00<br>120.00 | 90.00<br>90.00<br>120.00 | 90.00<br>90.00<br>90.00 | 90.00<br>90.00<br>120.00 | 90.00<br>90.00<br>90.00 | 90.00<br>90.00<br>120.00 | 90.00<br>90.00<br>90.00 |
| Mosaicity (°) | 0.05 | 0.05 | 0.08 | 0.07 | 0.06 | 0.06 | 0.06 | N/A | N/A | 0.45 | 0.61 |
| Resolution range (Å) | 37.75-1.05 (1.07-1.05) | 37.67-0.92 (0.94-0.92) | 39.24-0.94 (0.96-0.94) | 40.03-0.88 (0.89-0.88) | 37.72-1.00 (1.02-1.00) | 37.73-0.97 (0.99-0.97) | 40.04-0.98 (1.00-0.98) | 14.82-2.15 (2.23-2.15) | 14.61-1.79 (1.86-1.79) | 48.96-1.63 (1.66-1.63) | 100.18-1.65 (1.71-1.65) |
| Total no. of observations | 1215990 (57600) | 1844792 (80301) | 1624062 (71520) | 2138176 (92375) | 1469226 (68127) | 1599293 (71108) | 1498681 (42518) | 116016 (8779) | 224702 (11701) | 209375 (9255) | 382937 (23425) |
| No. of unique observations | 114819 (5635) | 168364 (8207) | 222540 (10804) | 269868 (13123) | 132397 (6477) | 144677 (7037) | 194272 (8354) | 12347 (1255) | 32647 (2863) | 31092 (1494) | 43186 (4304) |
| Completeness (%) | 100 (99.6) | 99.9 (99.1) | 99.9 (98.6) | 99.9 (99.1) | 100 (99.4) | 99.9 (99.0) | 99.0 (86.7) | 88.6 (92.4) | 97.1 (86.6) | 99.8 (99.7) | 99.7 (99.3) |
| Multiplicity | 10.6 (10.2) | 11.0 (9.8) | 7.3 (6.6) | 7.9 (7.0) | 11.1 (10.5) | 11.1 (10.1) | 7.7 (5.1) | 9.4 (7.0) | 6.9 (4.09) | 6.7 (6.2) | 8.9 (5.4) |
| ⟨I/σ(I)⟩ | 13.2 (0.6) | 18.2 (0.8) | 11.1 (0.6) | 16.0 (0.8) | 16.7 (0.8) | 16.4 (0.9) | 20.5 (2.0) | 9.7 (4.1) | 12.7 (3.1) | 21.4 (2.3) | 26.96 (1.59) |
| $CC_{1/2}$ | 0.999 (0.252) | 1.000 (0.332) | 0.999 (0.303) | 1.000 (0.395) | 1.000 (0.443) | 0.999 (0.394) | 1.000 (0.754) | 0.950 (0.490) | 0.905 (0.604) | 0.991 (0.833) | 0.995 (0.738) |
| $R_{meas}$ | 0.075 (3.838) | 0.057 (2.787) | 0.085 (3.572) | 0.060 (2.642) | 0.058 (3.036) | 0.065 (2.576) | 0.048 (0.790) | 0.287 (0.348) | 0.236 (0.305) | 0.049 (0.714) | 0.063 (0.589) |

**Table 2:** Crystallographic refinement statistics

| Sample | H-DJ-1 | D-DJ-1 | H-YajL | D-YajL | DJ-1 M17T | DJ-1 I21T | YajL D23N | DJ-1 neutron/X-ray joint refinement | YajL neutron/X-ray joint refinement |
|---|---|---|---|---|---|---|---|---|---|
| PDB code | 9YCU | 9YFR | 9YFS | 9YGW | 9YH8 | 9YGX | 9YKF | 9YZV | 9YZW |
| Refinement program | PHENIX 1.21.2-5419 | PHENIX 1.21.2-5419 | PHENIX 1.21.2-5419 | PHENIX 1.21.2-5419 | PHENIX 1.21.2-5419 | PHENIX 1.21.2-5419 | PHENIX 1.21.2-5419 | PHENIX 1.19.2_4158 | PHENIX 1.19.2_4158 |
| Resolution range (Å) | 37.55-1.05 (1.09-1.05) | 37.42-0.92 (0.95-0.92) | 39.24-0.94 (0.97-0.94) | 39.20-0.88 (0.91-0.88) | 37.52-1.00 (1.04-1.00) | 37.48-0.97 (1.00-0.97) | 39.17-0.98 (1.02-0.98) | Neutron:14.09-2.15 (2.36-2.15) X-ray: 33.13-1.63 (1.68-1.63) | Neutron:14.60-1.79 (1.85-1.79) X-ray:25.56-1.65 (1.69-1.65) |
| Completeness (%) | 99.52 (95.24) | 99.60 (96.91) | 99.21 (93.20) | 99.66 (97.68) | 99.69 (97.27) | 99.83 (98.40) | 98.92 (91.36) | Neutron: 88.03 (90) X-ray: 99.61 (99) | Neutron: 96.73 (81) X-ray: 99.64 (99) |
| No. of reflections | 114177 (10859) | 167834 (16178) | 220914 (20569) | 269187 (26146) | 131984 (12764) | 144423 (14106) | 194153 (17759) | Neutron: 12346 (3062) X-ray: 31054 (2747) | Neutron:32639 (2273) X-ray: 43084 (2619) |
| No. of reflections, test set | 3467 (289) | 5069 (458) | 6568 (633) | 8004 (776) | 4052 (390) | 4555 (508) | 5856 (589) | Neutron: 623 (153) X-ray: 1580 (141) | Neutron: 1659 (121) X-ray: 2215 (121) |
| $R_{work}$ | 0.1182 (0.3229) | 0.1039 (0.2837) | 0.1207 (0.3374) | 0.1123 (0.3087) | 0.1097 (0.2891) | 0.0994 (0.2673) | 0.0969 (0.1832) | Neutron: 0.2318 (0.3325) X-ray: 0.1334 (0.2269) | Neutron: 0.1774 (0.2939) X-ray: 0.1371 (0.2389) |
| $R_{free}$ | 0.1342 (0.3266) | 0.1125 (0.3019) | 0.1370 (0.3511) | 0.1258 (0.3282) | 0.1304 (0.2946) | 0.1127 (0.2531) | 0.1144 (0.1923) | Neutron: 0.2813 (0.3753) X-ray: 0.1548 (0.2250) | Neutron:0.2226 (0.3117) X-ray: 0.1642 (0.2715) |
| No. of non-H atoms | | | | | | | | | |
| Protein | 1583 | 1636 | 3375 | 3358 | 1564 | 1634 | 3611 | 1529 | 3094 |
| Water | 299 | 304 | 614 | 644 | 297 | 304 | 578 | 129 | 181 |
| Total | 1906 | 1948 | 3993 | 4006 | 1882 | 1958 | 4221 | 1658 | 3275 |
| Average R.M.S. deviations | | | | | | | | | |
| Bonds (Å) | 0.008 | 0.006 | 0.008 | 0.008 | 0.006 | 0.008 | 0.008 | 0.011 | 0.009 |
| Angles (°) | 1.08 | 1.01 | 1.08 | 1.17 | 1.02 | 1.15 | 1.14 | 1.46 | 1.22 |
| Average B factors (<$B_{iso}$>;Å$^2$) | | | | | | | | | |
| Protein | 16.85 | 12.49 | 11.61 | 11.01 | 14.86 | 13.98 | 9.72 | 41.69 | 24.55 |
| Water | 35.23 | 35.65 | 28.47 | 26.81 | 34.88 | 35.92 | 26.24 | 59.53 | 37.44 |
| Average ADP anisotropy[1] | | | | | | | | | |
| Protein | 0.563 | 0.546 | 0.414 | 0.391 | 0.587 | 0.567 | 0.490 | 0.560[2] | 0.564[2] |
| Water | 0.466 | 0.352 | 0.391 | 0.381 | 0.481 | 0.383 | 0.410 | 1 | 1 |
| MolProbity clashscore | 0.89 | 1.17 | 1.60 | 1.46 | 0.90 | 1.16 | 0.94 | 2.62 | 2.47 |
| Ramachandran plot | | | | | | | | | |
| Outliers (%) | 0.53 | 0.54 | 0.00 | 0.00 | 0.54 | 0.00 | 0.00 | 0.00 | 0.00 |
| Allowed (%) | 0.53 | 0.54 | 2.33 | 2.07 | 0.54 | 1.08 | 2.09 | 1.58 | 1.55 |
| Favored (%) | 98.93 | 98.92 | 97.67 | 97.93 | 98.92 | 98.92 | 97.91 | 98.42 | 98.45 |

[1]Anisotropy is defined as the ratio of the smallest to largest eigenvalue of the anisotropic ADP tensor.
[2]Refined using TLS for protein atoms and isotropic solvent in PHENIX.
